# Genetic code expansion enables plant-directed control of bacterial activity

**DOI:** 10.64898/2026.05.29.728790

**Authors:** Vivian Zhong, Michaela A. Jones, Avaniek Cabales, Alice Gevorgyan, René Inckemann, Anna A. Johnson, Sumudu S. Karunadasa, Amanda M. Forti, Shou-Ling Xu, Aditya M. Kunjapur, Jennifer A. N. Brophy

**Affiliations:** Department of Bioengineering, Stanford University, Stanford, CA 94305, USA; Department of Chemical and Biomolecular Engineering, University of Delaware, Newark 19716, DE, USA; Department of Biology, Stanford University, Stanford, CA 94305, USA; Department of Plant Biology and Carnegie Mass Spectrometry Facility, Carnegie Institution for Science, Stanford, CA 94305, USA; Howard Hughes Medical Institute, Stanford, CA 94305, USA

## Abstract

Programmable control of microbial gene expression by plant hosts could enable a new generation of precision agricultural biotechnology. Here, using *O*-methyl-L-tyrosine (OMY) as a model compound, we establish non-canonical amino acids (ncAAs) as a tool for plant-based control of associated microbial activity. We use genetic code expansion to engineer OMY-dependent control of protein synthesis in the soil bacterium *Bacillus subtilis.* Then, we engineer agronomically diverse plants, including Arabidopsis, tomato and poplar, to biosynthesize OMY. We show that plant-derived OMY can stimulate gene expression in both model and wild soil bacteria while also demonstrating that inducible and tissue-specific expression of a single biosynthetic enzyme by the plant enables on-demand control over microbial activity. This work establishes ncAAs as a tool for programming plant-microbe partnerships.

## Main Text

Agricultural productivity increased dramatically over the last century. Unfortunately, the fertilizers, growth regulators, and pesticides that drove these gains are broadly acting and often applied in excess, which has caused widespread waste and ecological damage^1,2^. Bacteria with the potential to deliver plant growth-promoting benefits directly to plants have emerged as promising alternatives to chemical inputs^3,4^. However, microbes may face analogous limitations to chemical inputs^5–7^. Beneficial microbes can spread beyond target crops, where they may confer growth benefits that dilute the competitive advantage intended for the crop and cause unintended ecological change^8,9^.

Solving this problem requires a mechanism to restrict microbial benefits to specific plant hosts and conditions. Previous efforts have explored rhizopines^10,11^, opines^12^, sucrose^13^, and quorum-sensing molecules^14,15^ as plant-produced signals to control bacterial gene expression. Though effective under the contexts tested, these molecules’ natural roles in inter-kingdom signaling could limit their orthogonality in synthetic signaling systems intended for sets of engineered organisms.

Non-canonical amino acids (ncAAs), a class of non-proteinogenic amino acids with diverse structures and functions, have potential for expanding the toolbox. Plants naturally produce an estimated 250-300 ncAAs^16–18^. These ncAAs serve as metabolic intermediates, protected nitrogen reservoirs, allelopathic chemicals, signals for systemic acquired resistance, and mediators of stress response^16–20^. Non-native ncAAs that are absent from natural plant chemical vocabularies could be combined with genetic code expansion (GCE) technologies to program plant-microbe interactions. GCE technologies now enable site-specific incorporation of more than 200 ncAAs into target proteins within living cells^21,22^, which creates the opportunity to program bacterial gene expression in response to diverse ncAAs.

Among the hundreds of ncAAs identified to date, *O*-methyl-L-tyrosine (OMY) is well suited for plant-based control over microbial behavior for several reasons. First, OMY has not been detected in agricultural soils and its known biosynthetic origins are limited to marine *Streptomyces* spp. and the rice blast fungus *Magnaporthe oryzae,* where it serves as a biosynthetic precursor for marformycin^23^ and cytochalasan^24^, respectively. Thus, OMY has a favorable likelihood of functioning as an orthogonal regulator of bacterial gene expression in agricultural soils. Second, OMY can be synthesized from L-tyrosine in a single enzymatic step by MfnG, an O-methyltransferase from *Streptomyces drozdowiczii*^23^, which facilitates its production in heterologous hosts^25^. Third, OMY was the first ncAA to be site-specifically incorporated within proteins in live bacterial cells^26^ and robust GCE machinery exists for OMY incorporation^27,28^. Finally, it has been demonstrated that OMY can mediate engineered communication between strains of *Escherichia coli*^25^.

Here, we establish OMY as a tool for plant-based control over microbial behavior. We demonstrate that the model soil bacterium *Bacillus subtilis* can incorporate OMY into target proteins through GCE to enable OMY-dependent protein synthesis with a >3000-fold dynamic range. We show that OMY can be biosynthesized by multiple plant species, including the model plant *Arabidopsis thaliana*, the vegetable crop *Solanum lycopersicum* (tomato), and the bioenergy tree *Populus tremula x alba* (poplar), through heterologous expression of a single biosynthetic enzyme. Plant-derived OMY is sufficient to control bacterial protein synthesis in co-cultivation experiments. We show that the system is generalizable across multiple target proteins and functional in a wild bacterial isolate with minimal optimization.

Translating this tool into agricultural applications requires that OMY production does not compromise plant fitness. Relative to plant-produced ncAAs that function as allelopathic compounds, such as meta-tyrosine^29^ and L-canavanine^30^, OMY exhibits reduced phytotoxicity. We find that OMY is ∼50-200-fold less potent than known allelopathic ncAAs and is not misincorporated into native plant proteins like other allelopathic ncAAs^16–19,29,31^. Conditional and tissue-specific production of OMY reduces growth impacts while maintaining effective bacterial signaling. Altogether, this work demonstrates that ncAAs can be used to program plant-microbe interactions and lays a foundation for precision agriculture applications based on engineered microbial partnerships.

## Results

### Genetic code expansion in *Bacillus subtilis*

To create bacteria capable of a highly sensitive response to plant-produced OMY (**Fig. 1a**), we introduced GCE components into *Bacillus subtilis* PY79^27,32^. GCE requires an orthogonal aminoacyl-tRNA synthetase/tRNA pair that can charge a suppressor tRNA with the ncAA to enable incorporation at amber stop codons in target genes. We introduced the napARS synthetase derived from the *Methanocaldococcus jannaschii* tyrosyl-tRNA synthetase (MjTyrRS) and its corresponding suppressor tRNA (tRNA^Tyr^_CUA_)^26^ under the control of strong constitutive promoters into the *B. subtilis* PY79 chromosome (**Fig. 1b**). To assess OMY-dependent protein production, we integrated an mNeonGreen reporter with an amber stop codon near the N-terminus at the *amyE* locus on the *B. subtilis* chromosome. In the absence of OMY, mNeonGreen translation should terminate at the stop codon, resulting in a truncated protein with no reporter activity. In the presence of OMY, napARS and tRNA^Tyr^_CUA_ should facilitate incorporation of OMY at UAG sites to produce a fully functional reporter protein. Testing of the mNeonGreen reporter by titrating exogenous OMY revealed dose-dependent reporter activation, with two-fold induction at 1 μM OMY, five-fold induction at 10 μM OMY, and 20-fold induction at 1 mM OMY (**Fig. 1c**). However, we observed substantial background fluorescence in the absence of OMY, indicating incomplete translational termination at the single amber stop codon.

**Figure 1.**
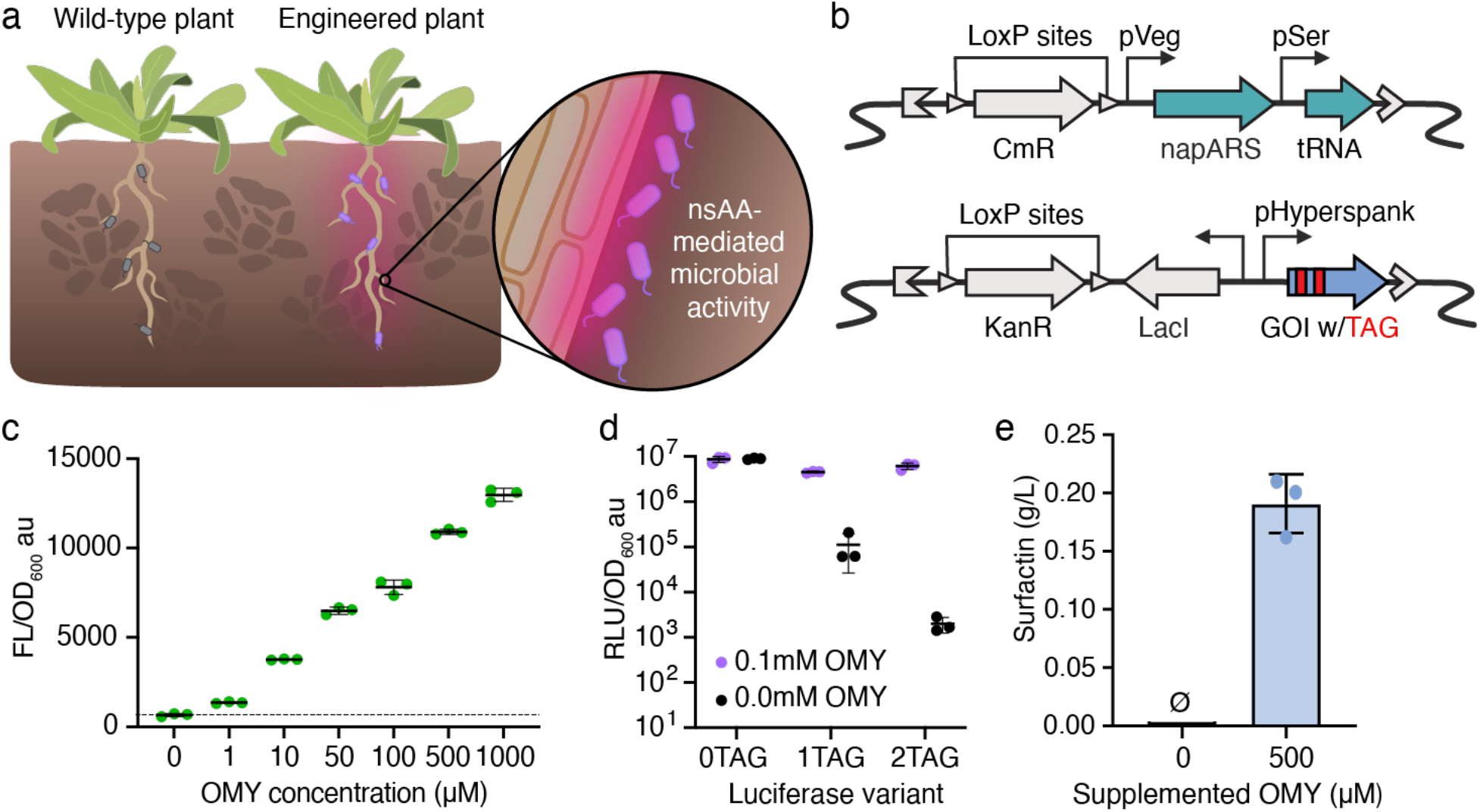
Genetic code expansion for O-methyl-L-tyrosine-dependent protein synthesis in a model soil bacterium. **(a)** OMY platform concept. Plants engineered to secrete a ncAA can activate specific bacterial activity. **(b)** Genetic code expansion components introduced to *B. subtilis* to enable OMY-dependent gene expression. An engineered variant of the *Methanocaldococcus jannaschii* tyrosyl-tRNA synthetase (MjTyrRS)^26^ and orthogonal suppressor tRNA (tRNA^Tyr^ _CUA_)^26^ were constitutively expressed by the pVeg^73^ and pSer ^27^ promoters, respectively. The Gene Of Interest, GOI, was engineered to have an in-frame amber stop codon, TAG, for site specific ncAA incorporation and was expressed constitutively by a pHyperspank promoter^74^. A chloramphenicol resistance cassette, CmR^75^, and kanamycin resistance cassette, KanR^76^, were used for selection of each construct upon genome integration and both were flanked by LoxP. **(c)** Activity of an OMY-dependent mNeonGreen reporter containing one amber stop codon in *B. subtilis* PY79 across a range of OMY concentrations in minimal media. The dashed line represents OD-normalized fluorescence with no supplemented OMY. **(d)** Activity of NanoLuc reporters containing zero, one, or two amber stop codons in *B. subtilis* PY79 with 0 µM (black) or 100 µM (purple) OMY supplementation in rich media (LB). **(e)** OMY-dependent surfactin production measured via high-performance liquid chromatography (HPLC). Data are the mean of three replicates collected simultaneously from three separate cultures. Error bars are s.d., and dots are individual data points.

To reduce background activity and improve dynamic range, we switched to the more sensitive NanoLuc luciferase reporter^33^ and tested the effect of incorporating multiple amber stop codons. We constructed NanoLuc variants with one (position 2) or two (positions 2 and 4) amber stop codons and measured luminescence in the presence and absence of OMY. While all variants showed similar luminescence signals in the presence of OMY, background activity decreased dramatically with the additional amber stop codon (**Fig. 1d**). The two-stop variant exhibited nearly 55-fold lower background than the single-stop construct, resulting in a total fold change of 3,100-fold with and without OMY (**Fig. 1d**). This highlights how variation in amber stop codons can alter the stringency of OMY-dependent protein production for conditional gene expression applications.

A key advantage of GCE is that bacterial output can be reprogrammed by introducing amber codons into any gene of interest without changing the tRNA/synthetase machinery. To demonstrate control over a microbially produced compound relevant to plant growth promotion, we engineered OMY-dependent surfactin biosynthesis by introducing a copy of the *sfp* gene, which encodes the 4’-phosphopantetheinyl transferase enzyme responsible for activating surfactin synthetases^34^, containing an amber stop codon into *B. subtilis* chromosome. The engineered strain produced no detectable surfactin in the absence of OMY and ∼0.18 g/L surfactin with 500 µM OMY (**Fig. 1e**). Together, these results show that OMY-dependent translation can be used to tightly control expression of diverse bacterial proteins.

### Plant biosynthesis of O-methyl-L-tyrosine (OMY)

Next, we engineered plants to produce OMY. We first tested OMY production in plant cells using transient expression of GFP-tagged MfnG (sfGFP-MfnG) in *Nicotiana benthamiana* leaves (**Fig. 2ab**). We used the *Arabidopsis thaliana UBQ10* promoter^35^ to drive expression of sfGFP-MfnG and *Agrobacterium tumefaciens* infiltration to deliver the construct into plant cells. At 3 days post-infiltration, leaves expressing the fusion construct were harvested for LC-MS analysis. Extracts from proUBQ10::sfGFP-MfnG leaf tissue contained a metabolite with the expected m/z for OMY and co-eluted with a commercial OMY standard, while no signal was detected in control samples (**Fig. 2b, Extended Data Fig. 1a**). Thus, OMY can be synthesized in plant cells by expressing MfnG.

**Figure 2.**
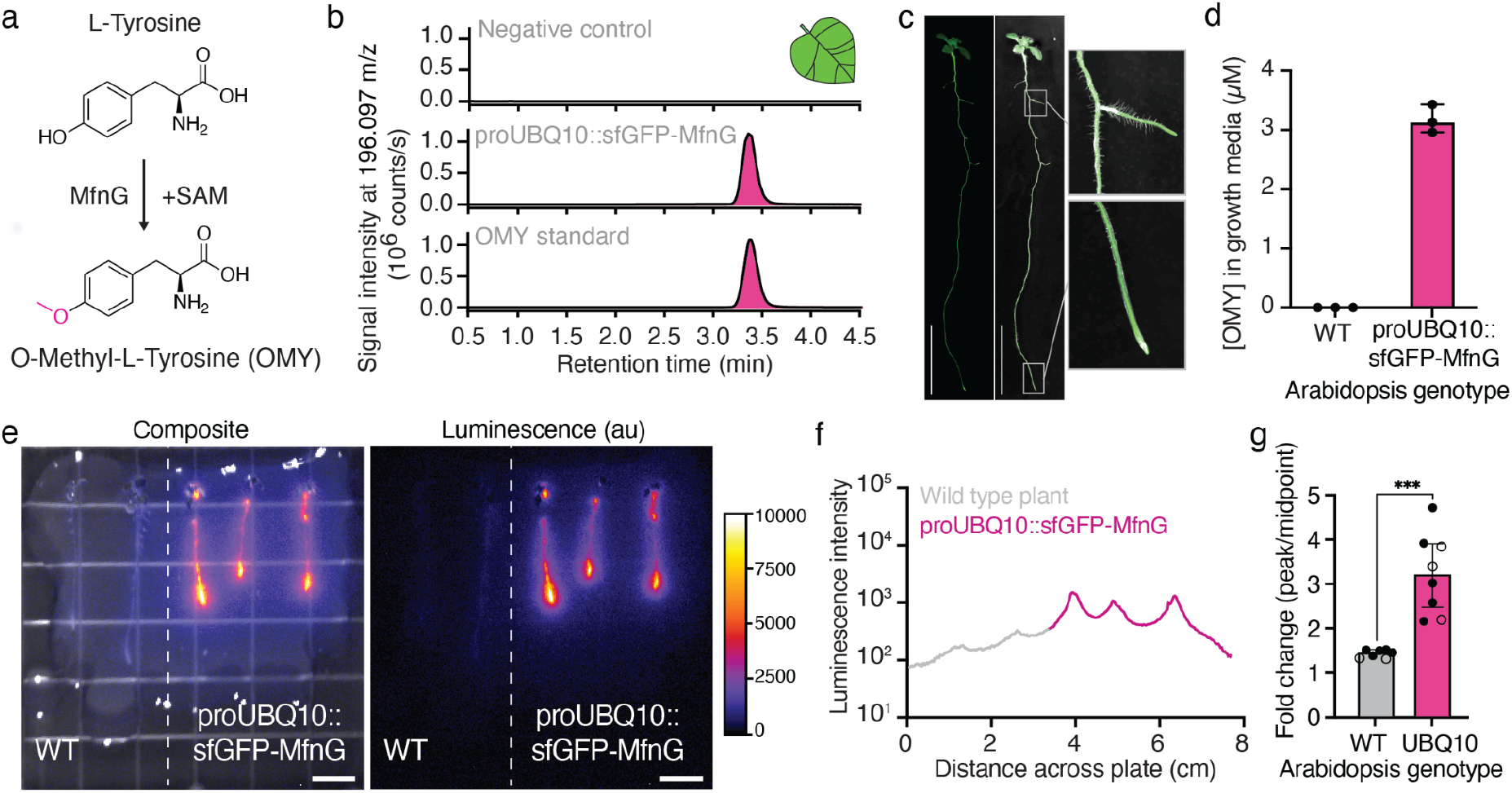
Heterologous MfnG expression enables OMY biosynthesis *in planta* and plant-directed control of bacterial gene expression. **(a)** O-methyl-L-tyrosine biosynthetic pathway. **(b)** Extracted ion chromatograms at 196.097 m/z from LC-MS analysis of uninfiltrated *Nicotiana benthamiana* leaves (top), *N. benthamiana* leaves transiently transformed with proUBQ10::sfGFP-MfnG T-DNA (middle), and OMY standard (bottom). Transient expression was achieved through Agrobacterium infiltration (Methods). **(c)** sfGFP expression in homozygous proUBQ10::sfGFP-MfnG seedlings 11 days post-stratification. Widefield fluorescence microscopy (500 ms exposure, left) and composite brightfield images (right). Insets show fluorescence of primary and lateral root tips. Scale bar, 1 cm. **(d)** OMY in growth media conditioned by hydroponically growing wild-type Col-0 (grey) and proUBQ10::sfGFP-MfnG (magenta) Arabidopsis seedlings. OMY concentration determined by peak intensity at 196.097 m/z from RT 4.855–5.139 min, EICs in Extended Data Fig. 1b. Dots represent three technical replicates from one batch of seedlings grown together, 18 seedlings per replicate. Error bars show the interquartile range. **(e)** Co-cultivation of *B. subtilis* PY79 expressing the OMY-dependent luciferase reporter containing two amber stop codons with wild-type Col-0 (left) or proUBQ10::sfGFP-MfnG (right) Arabidopsis seedlings. Seedlings were transferred onto bacterial-agar matrix at 8 days post-stratification and co-cultivated for 4 days before imaging. Composite images overlay brightfield with luminescence (30s exposure). Colorbar represents luminescence intensity (Fire LUT). Additional images in Extended Data Fig. 2. Scale bar, 1 cm. **(f)** Luminescence intensity profile across the x-axis of the plate in (e). Line plot connects discrete points representing the vertically averaged pixel intensity at each location. **(g)** Fold change in luminescence intensity, calculated as the ratio of intensity at each peak to the average intensity at adjacent midpoints between peaks. Bar plots show the median and interquartile range for wild-type Col-0 (grey, n=7) and proUBQ10::sfGFP-MfnG (magenta, n=8); individual data points represent single peaks. Open circles are from the mixed plate shown in (e), closed circles are from separate WT and proUBQ10::sfGFP-MfnG plates (Extended Data Fig. 2c). Statistical significance was evaluated using a Mann-Whitney test (***p < 0.001).

We then transformed *Arabidopsis thaliana* Col-0 with proUBQ10::sfGFP-MfnG via floral dip to generate stable transgenic lines, hypothesizing that biosynthesized OMY could be exuded because tyrosine and many other amino acids are commonly found in root exudate^36^. Transgenic plants exhibited sfGFP expression in all tissues (**Fig. 2c**). To quantify OMY exudation, we grew the plants hydroponically. We germinated Arabidopsis seeds on solid media and transferred 18 6-day old seedlings into 12 mL of liquid MS media (Murashige & Skoog basal salts) supplemented with 1% sucrose. After 48 hours of hydroponic growth, we analyzed the conditioned media by LC-MS. The proUBQ10::sfGFP-MfnG transgenic plants produced an average of 3.1 µM OMY, while no OMY was detected in wild-type controls (**Fig. 2d, Extended Data Fig. 1b**). Thus, micromolar OMY concentrations can be achieved in bulk root exudate without optimization of metabolic flux or OMY transport, and without L-tyrosine supplementation.

### Co-cultivation of engineered plants and bacteria

To test whether plant-produced OMY could control bacterial protein synthesis, we co-cultivated OMY-producing Arabidopsis with *B. subtilis* expressing OMY-dependent or OMY-independent NanoLuc reporters. We embedded the bacteria in MS agar media and transferred *Arabidopsis* seedlings onto the bacteria-agar matrix for co-cultivation (**Methods, Supplementary Fig. 1**). After 4-days of co-cultivation, we applied luciferase substrate and imaged luminescence. The OMY-independent reporter exhibited strong luminescence across the entire plate regardless of plant genotype, while OMY-dependent luciferase activity was only observed near sfGFP-MfnG expressing plants (**Fig. 2e, Extended Data Fig. 2a-c**). Plants expressing proUBQ10::sfGFP-MfnG produced strong luminescence peaks with the OMY-dependent reporter (∼3.2-fold above midpoint intensity) (**Fig. 2f-g, Extended Data Fig. 2c**). We also observed small luminescence peaks with the OMY-independent reporter at each seedling (∼1.2-fold above midpoint intensity), suggesting bacterial density may increase near the roots and contribute to a local increase in luminescence signal (**Extended Data Fig. 2a**). Wild-type plants co-cultivated with the OMY-dependent strain showed a similar, 1.4-fold increase in luminescence at the root (**Fig. 2f-g, Extended Data Fig. 2c**). Excitingly, when OMY-producing transgenic plants were grown side-by-side with unmodified wild-type plants, the luminescence signal was concentrated around the transgenic plants (**Fig. 2e-g**). Thus, plant-produced OMY enables genotype-specific induction of bacterial gene expression.

### OMY effects on plant and bacterial growth

Some plant-produced ncAAs function as allelopathic compounds that suppress the growth of competitor plants^16–20^. We noticed that transgenic plants ubiquitously expressing sfGFP-MfnG displayed reduced primary root length, decreased lateral root density, and impaired gravitropism compared to wild-type Col-0 (**Fig. 3a-d**). These phenotypes were consistent across multiple independent homozygous lines, suggesting that the growth defects result from sfGFP-MfnG overexpression rather than random T-DNA insertion effects (**Extended Data Fig. 3a, b**). Addition of OMY to bacterial growth media did not affect bacterial growth rate (**Extended Data Fig. 4a,b**).

**Figure 3.**
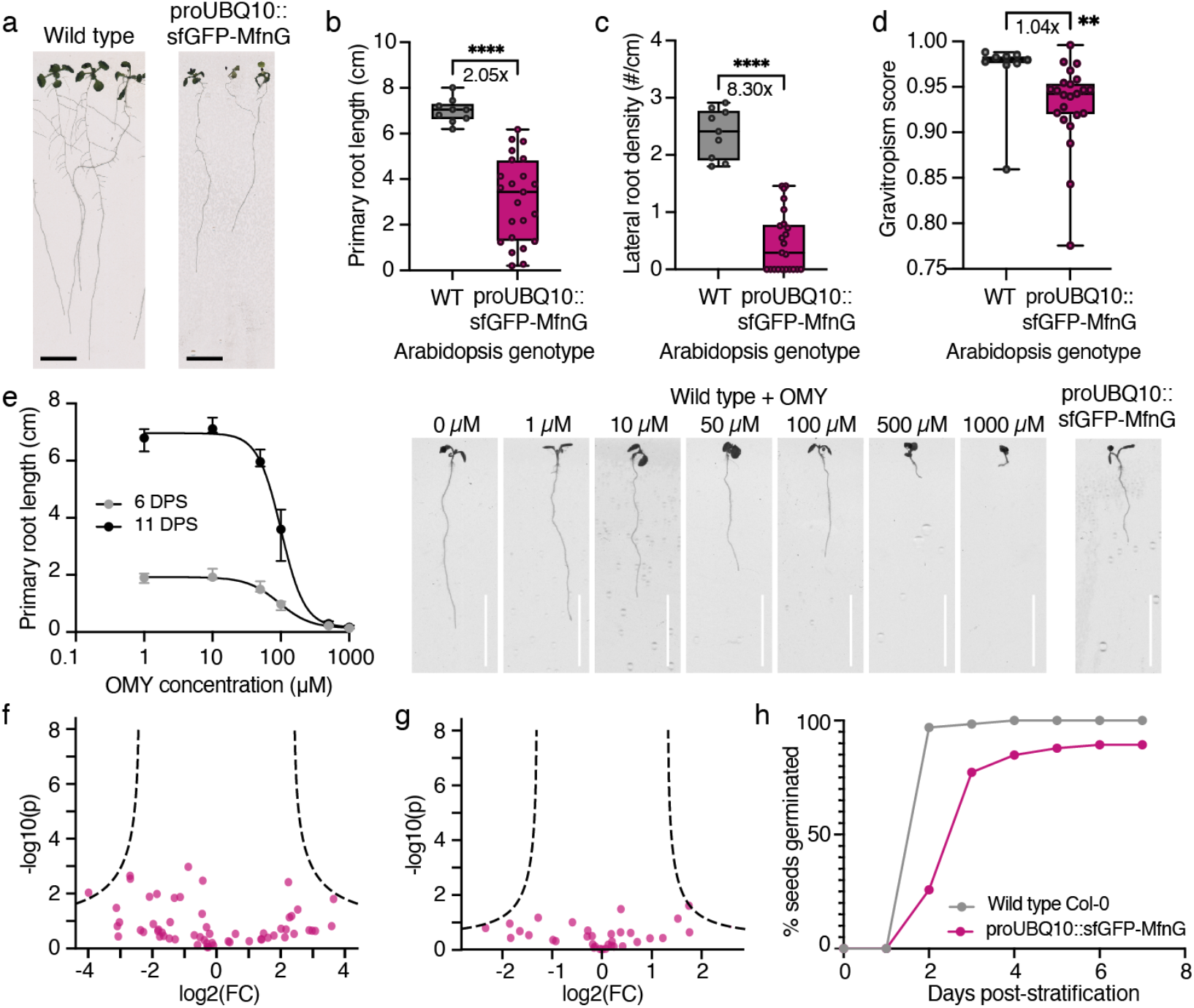
Characterization of OMY toxicity in Arabidopsis. **(a)** Representative images of wild-type Col-0 (left) and proUBQ10::sfGFP-MfnG (right) Arabidopsis seedlings 11 days post-stratification. **(b-d)** Primary root length (b), lateral root density (c), and gravitropism score (d) of wild-type Col-0 (grey, n=9) and proUBQ10::sfGFP-MfnG (magenta, n=23) Arabidopsis plants phenotyped at 11 days post-stratification. Box plots show the median, hinges indicate the first and third quartiles. Mann-Whitney tests were used to evaluate statistical significance, **** p < 0.0001, **p < 0.01. Primary root length measured from root-hypocotyl junction to the root tip. Lateral root density calculated as the number of emerged lateral roots divided by total primary root length. Gravitropism score calculated as the ratio of the vector length to primary root length. **(e)** Dose response curves for OMY inhibition of primary root length in wild-type Col-0 seedlings at 6 (grey) and 11 (black) days post-stratification. Representative images of seedlings at 6 days post-stratification. Dots are the median of 9-10 seedlings per concentration, error bars are IQR. IC50 curves fitted with Prism (Methods), R^2^>0.99 for both curves. **(f-g)** Volcano plots showing differential abundance of peptides containing O-methylated tyrosines (+14.0156 Da) in WT vs proUBQ10::sfGFP-MfnG shoots (f) and roots (g) harvested at 14 days post-stratification (n=3). Each dot is a unique peptide. Fold change (FC) is calculated as mean abundance in proUBQ10::sfGFP-MfnG samples divided by mean abundance in WT samples. Hyperbolic curves show thresholds for statistical significance determined with modified Student’s t-tests (curvature parameter (S_0_) = 1.5, FDR = 0.05). **(h)** Germination of wild-type Col-0 (grey) and proUBQ10::sfGFP-MfnG (magenta) Arabidopsis seeds over time. Unless otherwise stated, all panels show the same homozygous T3 proUBQ10::sfGFP-MfnG line (event #1). Scale bars, 1 cm.

To assess potential phytotoxicity of OMY, we treated wild-type Arabidopsis seedlings with OMY and measured primary root length. We observed concentration-dependent growth impairment and determined an IC50 of ∼94-101 µM (**Fig. 3e**). This makes OMY ∼50-200-fold less potent than known allelopathic ncAAs such as meta-tyrosine (IC50 2.4 µM)^29^ and L-canavanine (IC50 ∼0.5 µM)^30^. The concentration of OMY required to match the growth phenotype of proUBQ10::sfGFP-MfnG plants was ∼100-127 µM, which is consistent with constitutive overexpression driving substantial OMY accumulation *in planta*.

We investigated potential mechanisms underlying OMY-mediated growth inhibition. First, we tested whether inhibition resulted from depletion of aromatic amino acid precursors, which could occur through feedback inhibition of shikimate pathway enzymes^37^. However, supplementation with L-tyrosine (**Extended Data Fig. 5a**) or L-phenylalanine (**Extended Data Fig. 5b**) failed to rescue the growth phenotypes of OMY-producing plants. Next, we examined whether OMY phytotoxicity results from misincorporation into native proteins, which is a common mechanism of ncAA-dependent allelopathy^16–19,29,31^. Proteomics analysis did not detect significantly elevated levels of O-methylated L-tyrosine residues in proUBQ10::sfGFP-MfnG transgenic plants compared to wild-type (**Fig. 3f-g, Extended Data Fig. 5c, d**). Thus, we did not find evidence to support OMY misincorporation as the mechanism for toxicity.

Additional OMY phenotypes support a potentially novel mechanism of ncAA toxicity. proUBQ10::sfGFP-MfnG seeds exhibited delayed germination and reduced final germination percentages compared to wild-type (**Fig. 3h)**. proUBQ10::sfGFP-MfnG seed viability also declined more rapidly than Col-0, dropping from 90% germination at 3 months to 60% at 14 months (**Extended Data Fig. 5e**). Yet, exogenous OMY treatment of wild-type seeds had minimal effect on germination (**Extended Data Fig. 5f**). These germination phenotypes suggest OMY accumulation during seed development contributes to fitness costs in constitutively expressing lines.

### Conditional biosynthesis of O-methyl-L-tyrosine (OMY) *in planta*

To mitigate fitness costs while maintaining bacterial control, we engineered tissue-specific and inducible expression of MfnG. We selected the Arabidopsis *BEARSKIN1* (*BRN1*) promoter^38^ for root cap-specific expression, hypothesizing that this would limit systemic OMY accumulation while preserving rhizosphere accumulation. Transgenic Arabidopsis expressing proBRN1::sfGFP-MfnG displayed sfGFP signal exclusively in the root caps (**Fig. 4a**). In parallel, we engineered β-estradiol-inducible expression using the XVE system^39^. proXVE::sfGFP-MfnG plants showed no detectable sfGFP before treatment, but strong expression 24 hours after β-estradiol application (**Fig. 4b**).

**Figure 4.**
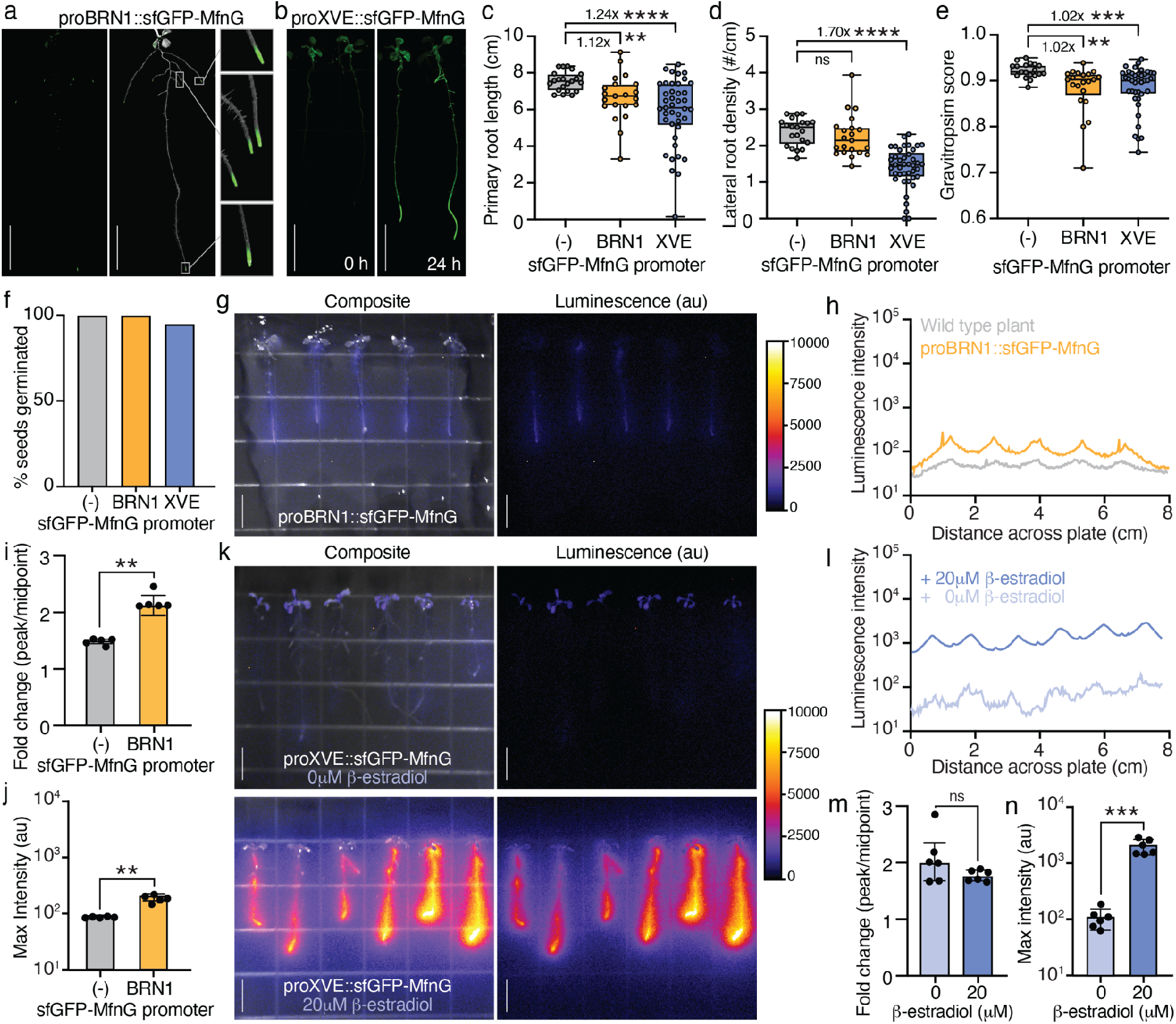
Conditional MfnG expression reduces fitness costs while maintaining plant-based bacterial control. **(a)** sfGFP expression in homozygous proBRN1::sfGFP-MfnG seedlings 11 days post-stratification. Widefield fluorescence microscopy (left, 500 ms exposure) and composite brightfield images (right). Insets show fluorescence of primary and lateral root tips. **(b)** sfGFP expression in T1 hemizygous transformants of proXVE::sfGFP-MfnG seedlings at 0 hours (left) and 24 hours (right) after transfer to media containing 20 µM ꞵ-estradiol. 150 ms exposure in the GFP channel. **(c-e)** Primary root length (c), lateral root density (d), and gravitropism score (e) of wild-type Col-0 (grey, n=21), proBRN1::sfGFP-MfnG (yellow, n=21), and proXVE::sfGFP-MfnG (blue, n=41) Arabidopsis plants, phenotyped 11 days post-stratification. Box plots show the median, hinges indicate the first and third quartiles. Statistical significance was evaluated using Kruskal-Wallis tests with Dunn’s post-hoc analysis (****p<0.0001, **p<0.01). (**f)** Germination of wild-type Col-0 (grey), proBRN1::sfGFP-MfnG (yellow), and proXVE::sfGFP-MfnG Arabidopsis seeds by 6 days post-stratification. **(g)** Co-cultivation of OMY-dependent *B. subtilis* PY79 luciferase reporter strain (as in Fig. 2e) with proBRN1::sfGFP-MfnG seedlings. Scale bars, 1 cm. **(h)** Luminescence intensity profile across the x-axis of the proBRN1::sfGFP-MfnG plate in (g) (orange) and wild-type Col-0 plate in Extended Data Fig. 2c (grey). **(i,j)** Fold change (i) and maximum (j) luminescence intensity for wild-type Col-0 (grey, n=5) and proBRN1::sfGFP-MfnG (orange, n=5) samples in (h). Statistical significance was evaluated using a Welch’s test (**p < 0.01). **(k)** Co-cultivation of OMY-dependent *B. subtilis* PY79 luciferase reporter strain (as in Fig. 2e) with proXVE::sfGFP-MfnG seedlings (event #1, right three; event #2, left three). 0 µM (top) or 20 µM (bottom) ꞵ-estradiol was added to the bacteria-agar matrix to induce sfGFP-MfnG expression. Scale bars, 1 cm. **(l)** Luminescence intensity profile across the x-axis of the proXVE::sfGFP-MfnG plates in (k). **(m,n)** Fold change (m) and maximum (n) luminescence intensity for proXVE::sfGFP-MfnG with 0 µM (light blue) or 20 µM (dark blue) ꞵ-estradiol. Statistical significance was evaluated using a Welch’s test (***p < 0.001). All panels utilize proBRN1::sfGFP-MfnG event #4 and proXVE::sfGFP-MfnG events #1 and #2.

Both proBRN1::sfGFP-MfnG and proXVE::sfGFP-MfnG plants exhibited substantially improved growth compared to constitutive proUBQ10::sfGFP-MfnG plants, with primary root length, lateral root density, and gravitropic response all closer to, or indistinguishable from, wild-type (**Fig. 4c-e**). Phenotypes varied minimally across independent T-DNA insertion lines (**Extended Data Figs. 3a, b**). Germination rates were comparable to wild-type at nearly 100% (**Fig. 4f**). Thus, conditional OMY production largely alleviates the fitness cost of constitutive MfnG overexpression.

We next tested whether conditional expression produces sufficient OMY for bacterial control. LC-MS analysis of conditioned media detected OMY in proBRN1::sfGFP-MfnG samples, though at lower concentrations than proUBQ10::sfGFP-MfnG (0.73 µM from 50 root cap-specific seedlings versus 3.1 µM from 18 ubiquitously expressing seedlings cultured in 12 mL media), which is consistent with *BRN1* expression being restricted to ∼19% of root cells^38,40^. Because this concentration falls below the initially characterized range (**Fig. 1c**), we tested incorporation at lower OMY concentrations. Using flow cytometry to sensitively measure OMY-dependent mNeonGreen fluorescence, we found induction at OMY concentrations as low as 0.05 µM (1.67-fold) and 0.5 µM (7.7-fold) (**Extended Data Fig. 4c**). Thus, the reporter remains sensitive to OMY below the 1 µM tested with plate reader fluorescence. In co-cultivation assays, proBRN1::sfGFP-MfnG plants activated OMY-dependent NanoLuc expression ∼2.2-fold above midpoint intensity (**Fig. 4g-j, Extended Data Fig. 2c**). Though luminescence was lower than observed with proUBQ10::sfGFP-MfnG plants, peaks at individual plant locations remained well defined.

Inducible expression of MfnG enabled strong induction of the bacterial reporter. We applied β-estradiol at the start of proXVE::sfGFP-MfnG plant co-cultivation with OMY-dependent bacteria by mixing it into the agar as it cooled. After four days of co-cultivation, induced plants (20 µM β-estradiol) showed a ∼18.7-fold increase in maximum luminescence intensity compared to uninduced plants (0 µM β-estradiol) (**Fig. 4k-n**). However, the fold change (peak/midpoint) in luminescence was not significantly different between induced and uninduced plants because luminescence did not decay to the same baseline observed in uninduced plants between peaks (**Fig. 4l-m**). Altogether, these results show that conditional OMY production maintains effective bacterial control while reducing growth inhibition.

### Engineering OMY-dependent gene expression in a plant growth-promoting bacterium

*B. subtilis* PY79 is a well-characterized model for Gram-positive soil bacteria, but its field application is limited. We therefore extended the OMY platform to *Bacillus subtilis* UD1022, a wild isolate that colonizes Arabidopsis roots and was previously reported to promote plant defense and growth^41^.

We first engineered *B. subtilis* UD1022 with the same GCE components used in the model strain *B. subtilis* PY79 and tested OMY-dependent mNeonGreen expression. UD1022 exhibited dose-dependent fluorescence similar to PY79 (**Fig. 5a, Extended Data Fig. 4d**). We next tested the activity of NanoLuc reporters containing varying numbers of amber stop codons. While the single-amber variant showed robust OMY-dependent luminescence in UD1022, the two-amber variant only reduced background by 4-fold and decreased maximal signal. Consequently, the optimal fold change for UD1022 (111-fold) was achieved with one amber codon, which differs from PY79 that demonstrated maximum luminescence fold change with two amber codons (**Fig. 5b**)

**Figure 5.**
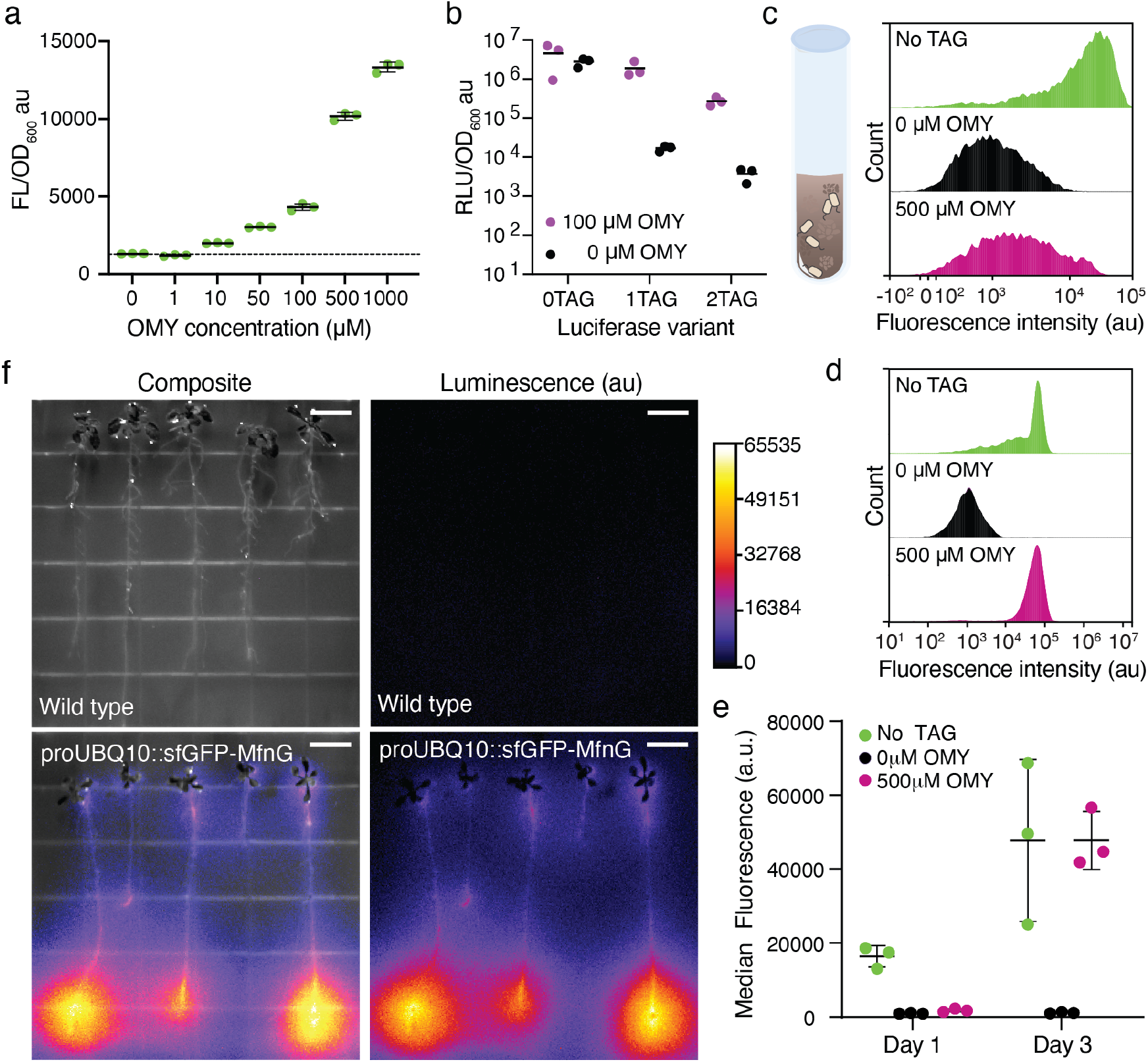
OMY-dependent gene expression in a plant growth-promoting bacterium. **(a)** Activity of an OMY-dependent mNeonGreen reporter in *Bacillus subtilis* UD1022 across a range of OMY concentrations in minimal media. **(b)** Activity of NanoLuc reporters containing zero, one, or two amber stop codons in *B. subtilis* UD1022 with 0 µM (black) or 100 µM (purple) OMY supplementation in rich media (LB). Data are the mean of three replicates collected simultaneously from three separate cultures. Error bars show s.d. and dots are individual data points. **(c-d)** Representative flow cytometry histograms of 10,000 events of UD1022 strains harboring a OMY-independent mNeonGreen reporter (green) or OMY-dependent mNeonGreen reporter extracted from autoclaved soil supplemented with 0 µM OMY (black) or 500 µM OMY (pink). Cells were extracted from the soil 1 day (c) or 3 days (d) after inoculation. **(e)** Median fluorescence for the same strains and conditions as (c-d) at 1 day and 3 days after inoculation for three replicates. **(f)** Co-cultivation of *B. subtilis* UD1022 expressing the OMY-dependent luciferase reporter containing two amber stop codons with wild-type Col-0 (top) or proUBQ10::sfGFP-MfnG (bottom) Arabidopsis seedlings. Composite images overlay luminescence (10s exposure) and brightfield. Seedlings transferred to bacterial-agar matrix at 10 days post-stratification and co-cultivated for 4 days. Colorbar represents luminescence intensity (Fire LUT). Additional images in Extended Data Fig. 6. Scale bars, 1 cm.

To test OMY-dependent gene expression in a more field-relevant environment, we inoculated autoclaved soil with the UD1022 mNeonGreen (1UAG) reporter strain and supplemented the soil with 500 μM OMY (**Extended Data Fig. 6a**). After 1-3 days, we quantified mNeonGreen expression by washing the soil with PBS, filtering out soil particles, and quantifying fluorescence by flow cytometry. Median fluorescence increased 41-fold between 0 μM and 500 μM OMY treatment at day 3, indicating robust OMY incorporation in soil (**Fig. 5c-e**).

Next, we tested whether plant-produced OMY could activate the UD1022 reporter in soil. We grew proUBQ10::sfGFP-MfnG or wild-type seedlings in autoclaved soil for 11 days, then inoculated the soil with UD1022 expressing the OMY-dependent mNeonGreen reporter. After three days of co-cultivation, we extracted bacteria for flow cytometry analysis. Bacteria containing the OMY-dependent mNeonGreen reporter showed elevated median fluorescence (802 a.f.u.) when co-cultivated with proUBQ10::sfGFP-MfnG plants compared to wild-type controls (530 a.f.u.). The total fluorescent population also increased from 9.8% to 36.1%, demonstrating plant-to-microbe OMY transfer in soil (**Extended Data Fig. 6b-d**).

To visualize spatial patterns of bacterial activity, we co-cultivated UD1022 expressing the OMY-dependent NanoLuc reporter with proUBQ10::sfGFP-MfnG Arabidopsis on agar. UD1022 displayed luminescence patterns similar to PY79, with signal brightest at the root tips despite ubiquitous MfnG expression (**Fig. 5f, Extended Data Fig. 7**). The root tip signal was more pronounced with UD1022, which may reflect its greater ability to utilize root exudates for growth. These results demonstrate that OMY-mediated control extends to wild soil bacteria with minimal optimization and functions in both agar and soil environments.

### Introducing OMY production to crop species

To demonstrate agricultural applicability, we extended OMY biosynthesis to tomato (*Solanum lycopersicum* cv. Heinz 1706^42^) and poplar (*Populus tremula x alba* INRA 717-1B4^43^). Since generating stable transgenics in these species is time-consuming, we used *Rhizobium rhizogenes*-mediated hairy root transformation for rapid proof-of-concept. In this process, *R. rhizogenes* transfers T-DNA into plant cells at wound sites, triggering the formation of transgenic adventitious roots^44^ **(Fig. 6a**). The resulting composite plants contain wild-type shoots and a mixture of transgenic hairy roots and non-transgenic roots emerging from the shoot^45^ (**Extended Data Fig. 8**).

**Figure 6.**
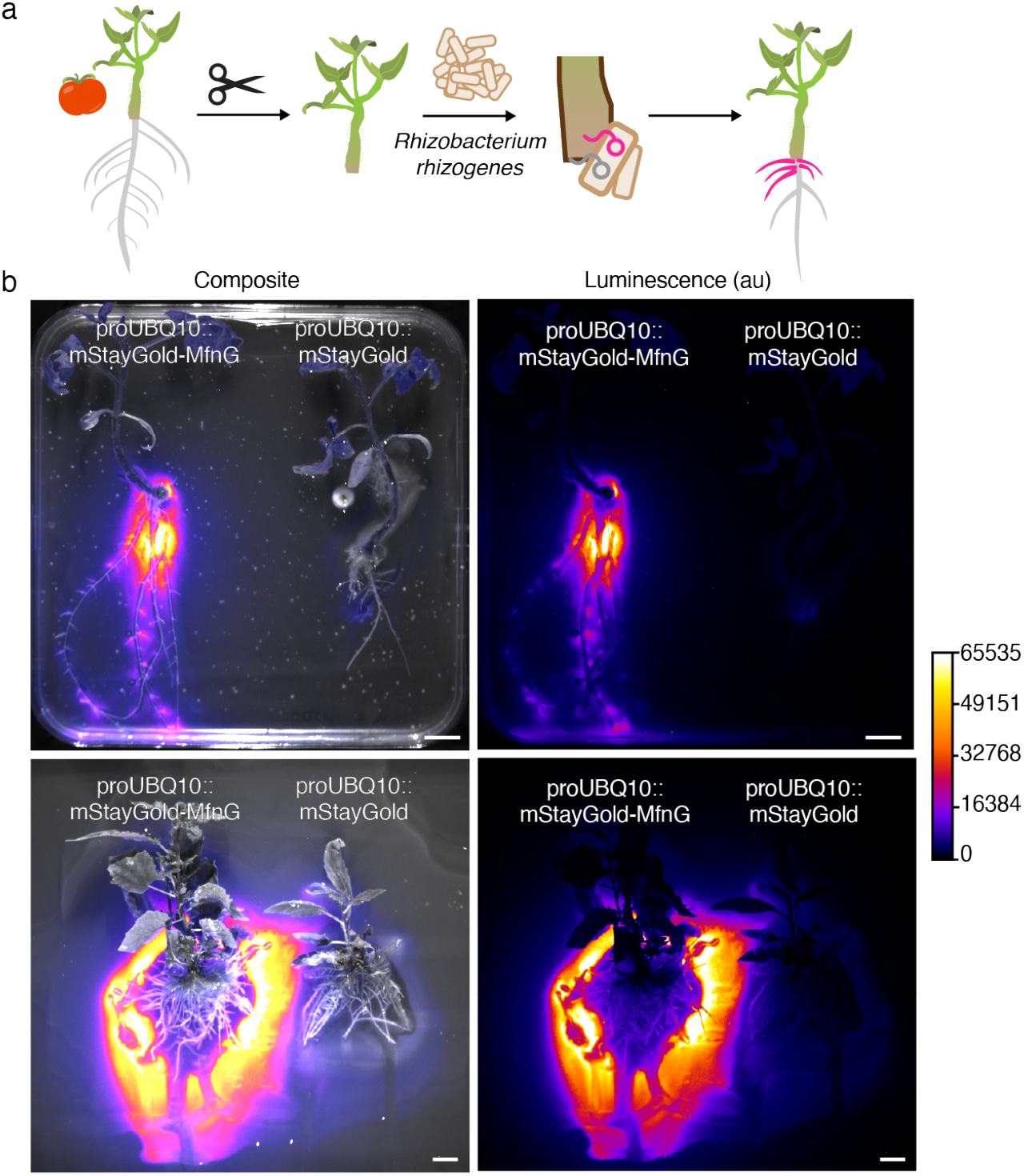
OMY biosynthesis in tomato and poplar drives genotype-specific bacterial reporter expression. **(a)** Hairy root transformation method for tomato composite plants. Roots were removed from 8-day old seedlings and inoculated with *Rhizobium rhizogenes* carrying the genes of interest. *R. rhizogenes* transfers its T-DNA into tomato cells, which induces hairy root growth and produces composite plants with wild-type shoots and a mixture of transgenic (magenta) and non-transgenic (grey) roots. **(b)** Co-cultivation of composite tomato (top) and poplar (bottom) plants with *B. subtilis* PY79 expressing an OMY-dependent luciferase reporter. Composite images overlay luminescence (30s exposure) and brightfield. Images collected after 5 days (tomato) and 4 days (poplar) of co-cultivation with bacteria. Colorbar represents luminescence intensity (Fire LUT). Scale bars, 1 cm.

We transformed tomato seedlings and poplar explants with proUBQ10::mStaygold-MfnG constructs (**Extended Data Fig. 8**). Co-cultivation of composite plants with PY79 expressing the OMY-dependent NanoLuc reporter for 4-5 days resulted in luminescence localized to fluorescent marker-expressing transgenic roots in both species (**Fig. 6b**). These results demonstrate that OMY biosynthesis and bacterial control are extensible to agriculturally relevant crop species.

## Discussion

Our work establishes the ncAA OMY as a tool for plant-based control of microbial activity. By combining OMY biosynthesis in plants with bacterial GCE, we empower plants to direct bacterial gene expression. The system operates with high specificity, which is visible in co-cultivation experiments where the luminescence signal from engineered bacteria concentrates around transgenic plants growing alongside wild-type neighbors. Though we find that OMY exhibits mild phytotoxicity, this occurs through a mechanism distinct from known allelopathic ncAAs^29,30^, and conditional OMY biosynthesis can ameliorate fitness costs while maintaining bacterial control. Future work can explore additional hypotheses for OMY toxicity, including competitive inhibition of aromatic amino acid transporters and biosynthetic enzymes^46^, interference with auxin biosynthesis or signaling (consistent with impaired root gravitropism)^47^, and interference with gibberellic acid biosynthesis or signaling (consistent with reduced germination)^48^, as well as methods for amelioration.

Future work can also optimize the system for agricultural soils, where resident bacteria and environmental fluctuations may degrade OMY, influence incorporation, and make it difficult for engineered strains to compete in the rhizosphere. Greater ncAA sensitivity, achieved through engineering of the aminoacyl-tRNA synthetase/tRNA pair for greater translation efficiency at low ncAA concentrations^49^, may make signaling more robust to these fluctuations without requiring phytotoxic levels of OMY production. Engineering natural field isolates may help address colonization and competition challenges. However, we found that maximum fold-change required different amber codon configurations in PY79 and UD1022. This may reflect variation in amber suppression efficiency^27,50,51^, background translational termination fidelity^52,53^, or ncAA transport and availability^54,55^ between strains. Thus, field deployment may require optimization of the orthogonal translation system and the amber codon number in each bacteria with plant hosts in the target environment. Testing this system in non-sterile soils with diverse native microbial communities will also be important for evaluating potential off-target effects and confirming that OMY-based signaling remains specific to engineered strains in complex, multi-species environments.

Interestingly, bulk measurements of OMY did not reflect the level of reporter activation at plant roots in co-cultivation experiments. This is likely because bulk measurements underestimate OMY concentration at the root, where bacteria contact the plant directly^56,57^. Though OMY concentration gradients may explain the strong luminescence of OMY-dependent reporter strains grown with MfnG-expressing plants, the OMY-independent reporter also showed a modest increase in luminescence near the roots of every plant genotype, suggesting bacterial cell density may increase near the roots^58^ and contribute to a local boost in luminescence signal. Though we did not directly measure OMY concentration or bacterial cell density at the root, labeling bacteria with a constitutive fluorescent marker, quantifying OMY and bacterial CFU extracted from the gel matrix at different plate locations would help measure the contribution of chemical and cell concentration gradients to reporter signals in future work.

Here, we extend GCE technology from intracellular protein engineering to programmable interactions between organisms. Historically, GCE largely focused on engineering new chemistries within a single organism^59^, including exogenous supply of synthetic ncAAs to control gene expression via translational switches^27,60^ and survival of engineered microbes^61,62^. Pioneers in the field have long speculated that these kinds of orthogonal biological systems would improve biosafety of engineered organisms by enabling one kind of “genetic firewall” that would limit horizontal gene transfer^63–65^. Although these tools and principles each pertain to applications in environmental biotechnology, prior work has not realized the creation of a self-sufficient pair of engineered organisms sharing a private communication channel via GCE. One exception is our own recent work where we demonstrated a proof-of-concept of ecological control of gene expression and survival among two engineered *E. coli* strains^25^. The system that we present here represents proof-of-concept that GCE can link plants and environmentally relevant microbes across kingdoms using an ncAA as the shared signal, with autonomous and spatiotemporally controlled synthesis of the ncAA by a multicellular organism.

NcAA-enabled plant-based control of microbial activity has potential for diverse future applications. While we demonstrated control over reporter expression and surfactin biosynthesis, the same approach could be used to regulate the production of other plant growth-promoting proteins, such as biosynthetic enzymes for antipathogen compounds, metal chelators, and mineral-solubilizing enzymes^66,67^. Similarly, tissue-specific and inducible expression of MfnG could be extended to engineer systems where bacterial activity responds to plant perception of stress, nutrient status, or developmental cues^68^. Excitingly, ncAAs open the door to biosynthesis of proteins with non-natural chemistries in the rhizosphere^69,70^. By incorporating ncAAs into select proteins, engineered microbes may make entirely new molecules for plants. The overall concept we present here is also readily extensible to other ncAAs. Hundreds of ncAAs have been identified in nature^16,71^, and dozens more can be accessed via designed semi-synthetic pathways^72^. Development of mutually orthogonal tRNA-synthetase pairs could enable multiplexed signaling where different plant cues trigger distinct bacterial responses. Altogether, ncAAs represent a vast and largely untapped vocabulary for engineering plant control over their microbiomes.

## Extended Data Figures

**Extended Data Fig. 1.**
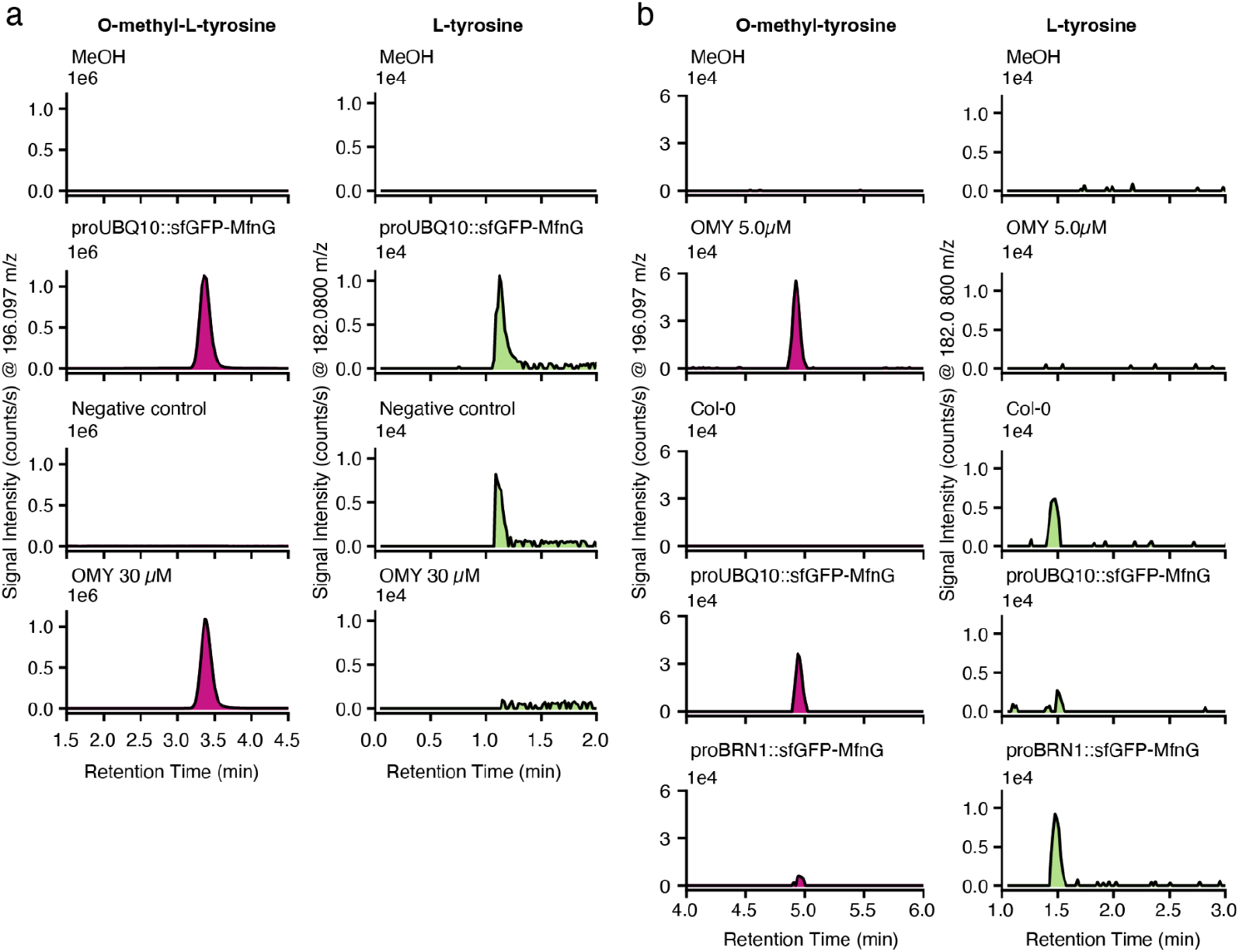
LC-MS detection of OMY and L-tyrosine in Nicotiana and Arabidopsis. **(a)** Representative extracted ion chromatography (EIC) traces at the m/z for OMY (left) and L-tyrosine (right) from LC-MS analysis of *Nicotiana benthamiana* leaf extracts. Lyophilized leaf disc samples and standards were prepared in 80% methanol, which served as the blank for the LC-MS run. Negative control was an uninfiltrated leaf. OMY 30 µM is the commercial standard. **(b)** Representative extracted ion chromatography (EIC) traces at the m/z for OMY (left) and L-tyrosine (right) from LC-MS analysis of metabolites extracted from conditioned hydroponic growth media. Lyophilized root exudate samples and standards were prepared in 80% methanol, which served as the blank for the LC-MS run. OMY 5 µM is the commercial standard.

**Extended Data Fig. 2.**
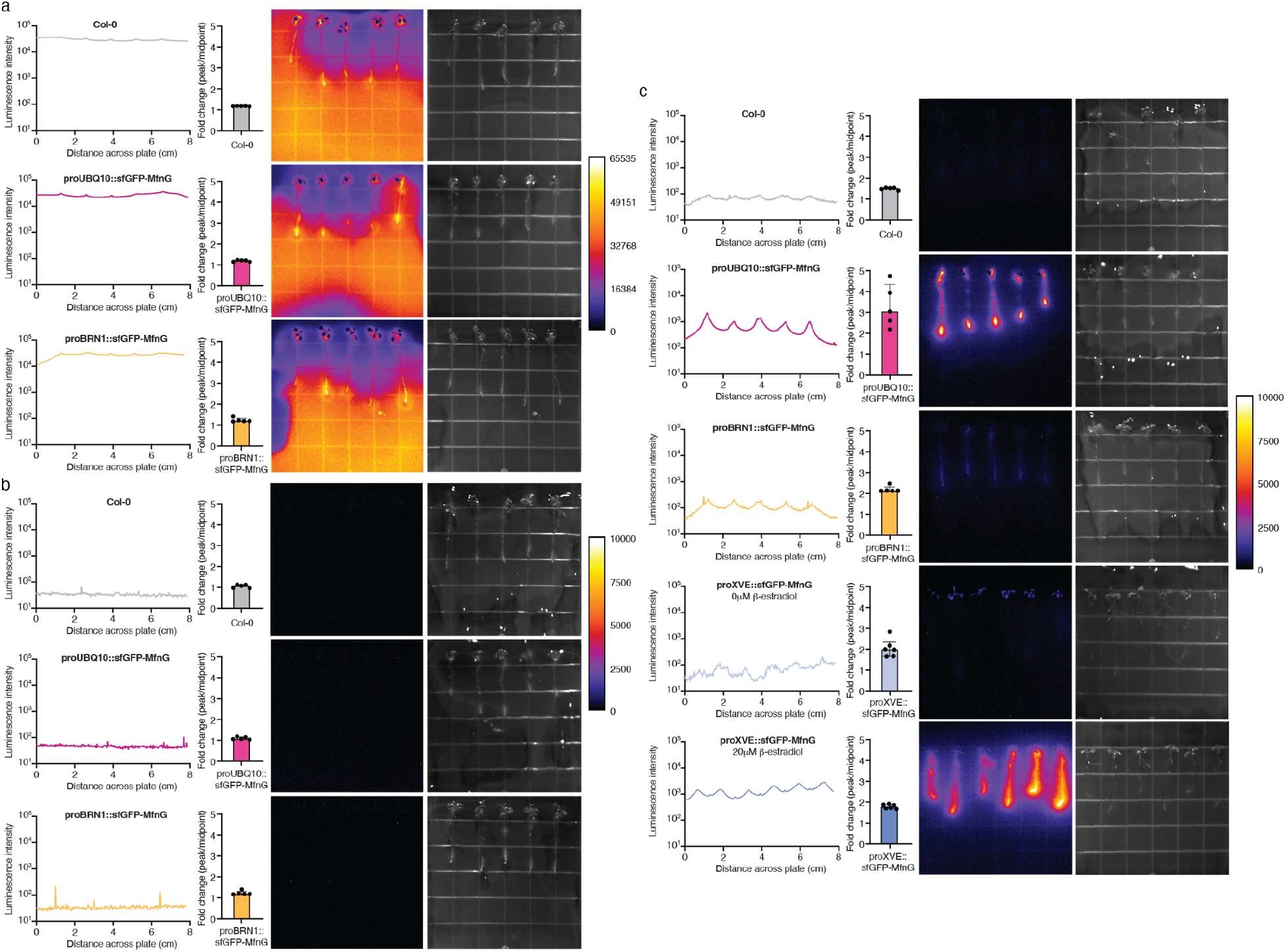
Additional controls for OMY-dependent luciferase expression in *B. subtilis* PY79 co-cultivated with Arabidopsis. **(a)** Co-cultivation of *B. subtilis* PY79 expressing the OMY-independent luciferase reporter with wild-type Col-0 (top, grey), proUBQ10::sfGFP-MfnG (middle, pink) or proBRN1::sfGFP-MfnG (bottom, orange) Arabidopsis seedlings. **(b)** Co-cultivation of wild-type *B. subtilis* PY79 with wild-type Col-0 (top, grey), proUBQ10::sfGFP-MfnG (middle, pink) or proBRN1::sfGFP-MfnG (bottom, orange) Arabidopsis seedlings. **(c)** Co-cultivation of *B. subtilis* PY79 expressing the OMY-independent luciferase reporter containing two stop codons with wild-type Col-0 (grey), proUBQ10::sfGFP-MfnG (pink), proBRN1::sfGFP-MfnG (orange), or proXVE::sfGFP-MfnG (blue) Arabidopsis seedlings. For proXVE::sfGFP-MfnG experiments, 0 µM (top) or 20 µM (bottom) ꞵ-estradiol was added to the bacteria-agar matrix to induce sfGFP-MfnG expression. In all panels, seedlings were transferred onto bacterial-agar matrix at 8 days post-stratification and co-cultivated for 4 days before imaging. Luminescence (30s exposure, left) and brightfield (right) images were collected with iBright imager. Colorbar represents luminescence intensity (Fire LUT). Line graphs show luminescence intensity profile across the x-axis of the co-cultivation plate; lines connect discrete points representing the vertically averaged pixel intensity at each location. Bar charts show fold change in luminescence intensity, calculated as the ratio of intensity at each peak to the average intensity at adjacent midpoints between peaks; individual data points represent single peaks. Panel (c) shows uncropped versions of images presented in Figs. 2 and 4.

**Extended Data Fig. 3.**
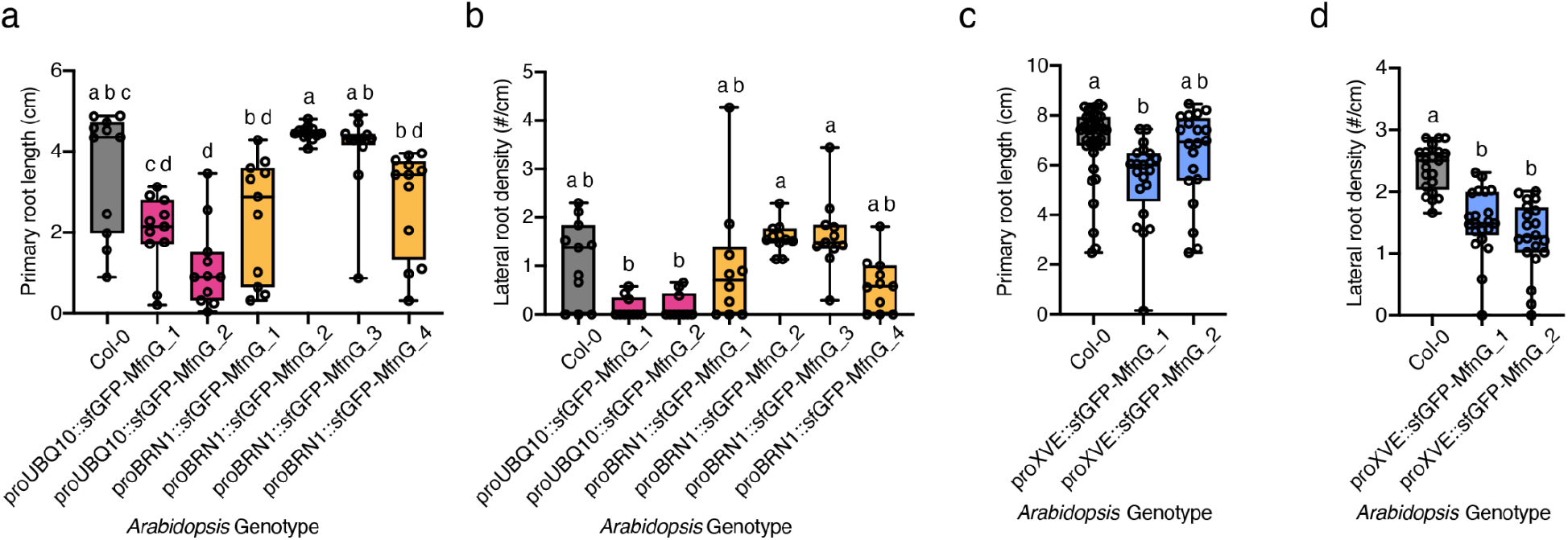
Consistency of growth phenotypes across independent T-DNA insertion lines. **(a-b)** Primary root length (a) and lateral root density (b) of independent homozygous proUBQ10::sfGFP-MfnG (magenta) and proBRN1::sfGFP-MfnG (yellow) lines T-DNA insertion lines. Box plots show the median of 11-12 plants phenotyped at 8 days post-stratification. Box plot hinges indicate the first and third quartiles. **(c-d)** Primary root length (c) and lateral root density (d) of independent T3 homozygous proXVE::sfGFP-MfnG lines. Box plots show the median of 21 plants phenotyped at 11 days post-stratification. Box plot hinges indicate the first and third quartiles. For all panels, significance groups were determined using Kruskal-Wallis tests with Dunn’s post-hoc analysis (*a*<0.05).

**Extended Data Fig. 4.**
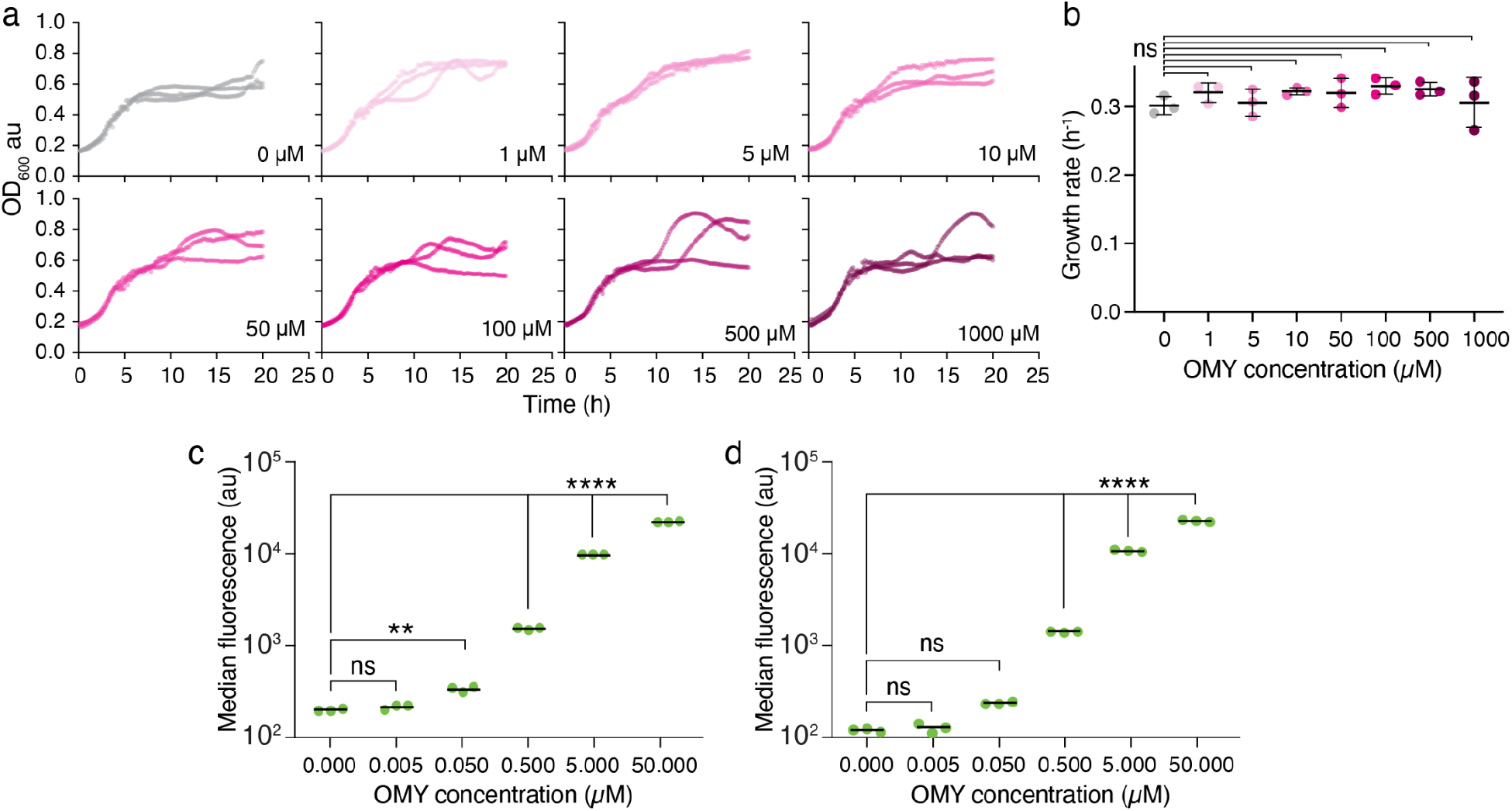
*B.subtilis* growth and protein expression with OMY. **(a)** Growth of PY79 expressing the amino-acyl tRNA synthetase and tRNA for ncAA incorporation in 1/2x MS media supplemented with 1.5% sucrose, 10 mM L-glutamate, and varying concentrations of OMY at 30°C. Each plot contains the growth for three individual biological replicates. The OMY concentration is indicated in the lower right corner of each plot. **(b)** Growth rates calculated for curves in (a). Data shows mean and standard deviation across three biological replicates. Statistical significance was determined using one-way ANOVA and Dunn’s test against the no OMY control (α = 0.05). **(c-d)** Fluorescence of *B. subtilis* PY79 (c) or UD1022 (d) expressing an OMY-dependent mNeonGreen reporter across a range of OMY concentrations in 1/2x MS media supplemented with 1.5% sucrose and 10 mM L-glutamate at 22°C. Fluorescence measured via flow cytometry. Statistical significance was evaluated using a one way ANOVA with Tukey HSD posthoc test (**p<0.01, ****p < 0.0001).

**Extended Data Fig. 5.**
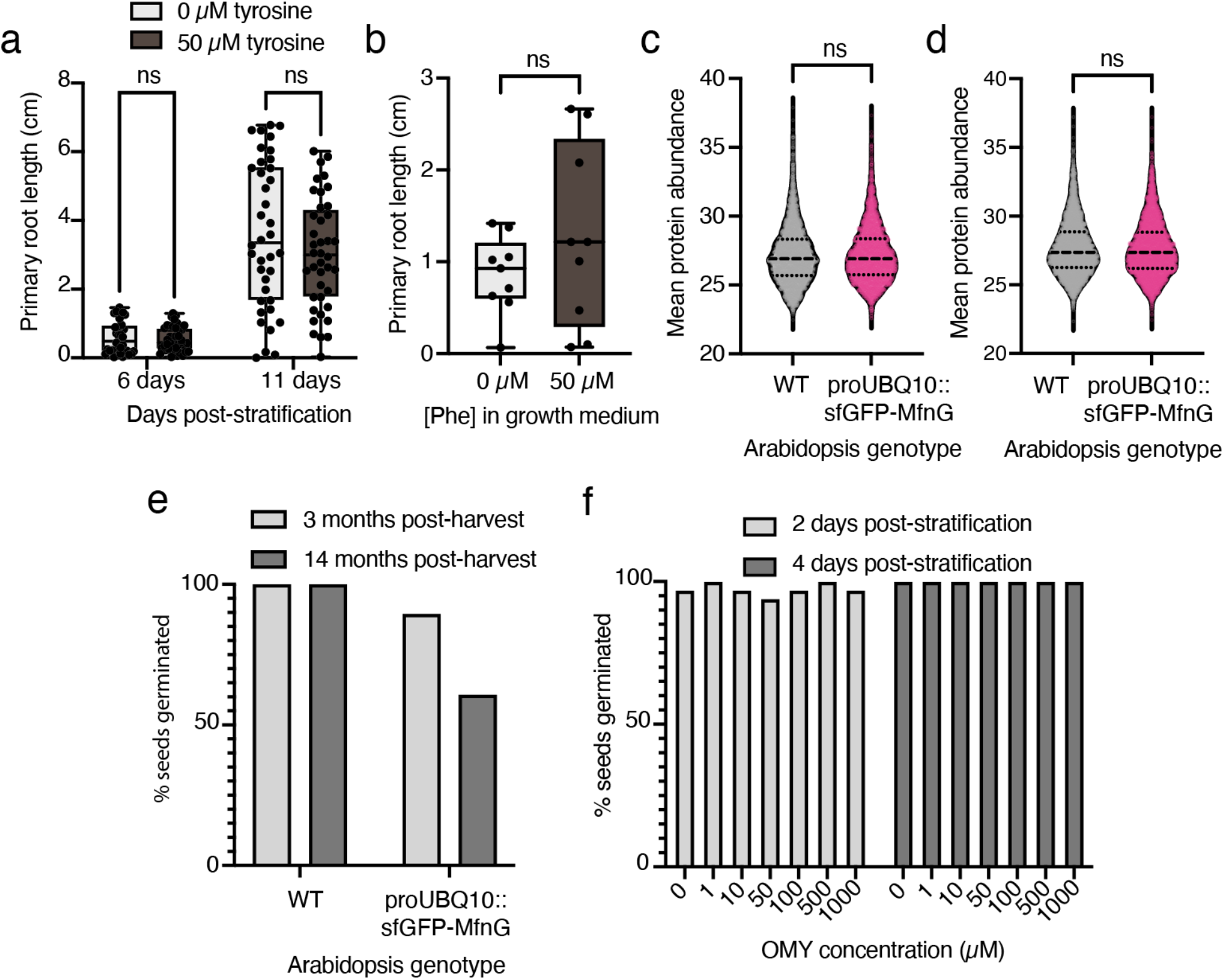
Investigation of OMY phytotoxicity mechanisms and germination phenotypes. **(a)** Primary root length of proUBQ10::sfGFP-MfnG seedlings supplemented with 0 µM or 50 µM l-tyrosine at 6 and 11 days post-stratification. Each datapoint represents an individual seedling; only germinated seeds were included. Statistical significance assessed by Mann-Whitney test. **(b)** Primary root length of proUBQ10::sfGFP-MfnG seedlings supplemented with 0 µM or 50 µM l-phenylalanine at 11 days post-stratification. Each datapoint represents an individual seedling; only germinated seeds were included. Statistical significance assessed by Mann-Whitney test. **(c-d)** Violin plots showing the mean abundance (n=3) for all detected protein species in WT and proUBQ10::sfGFP-MfnG shoot (c) and root (d) tissue, corresponding to Fig. 3f-g. Two-sided unpaired student’s t-test, p>0.05. Dashed lines mark the median and dotted lines mark quartiles. Dots are individual protein species. **(e)** Germination of wild-type Col-0 and proUBQ10::sfGFP-MfnG seeds at 3 months and 14 months post-harvest. Percent germination assessed at 4 days post-stratification. **(f)** Germination of wild-type Col-0 seeds at 2 days and 4 days post-stratification across a range of OMY concentrations. One-tailed Fisher’s exact test used to evaluate statistical significance (p=1e-4).

**Extended Data Fig. 6.**
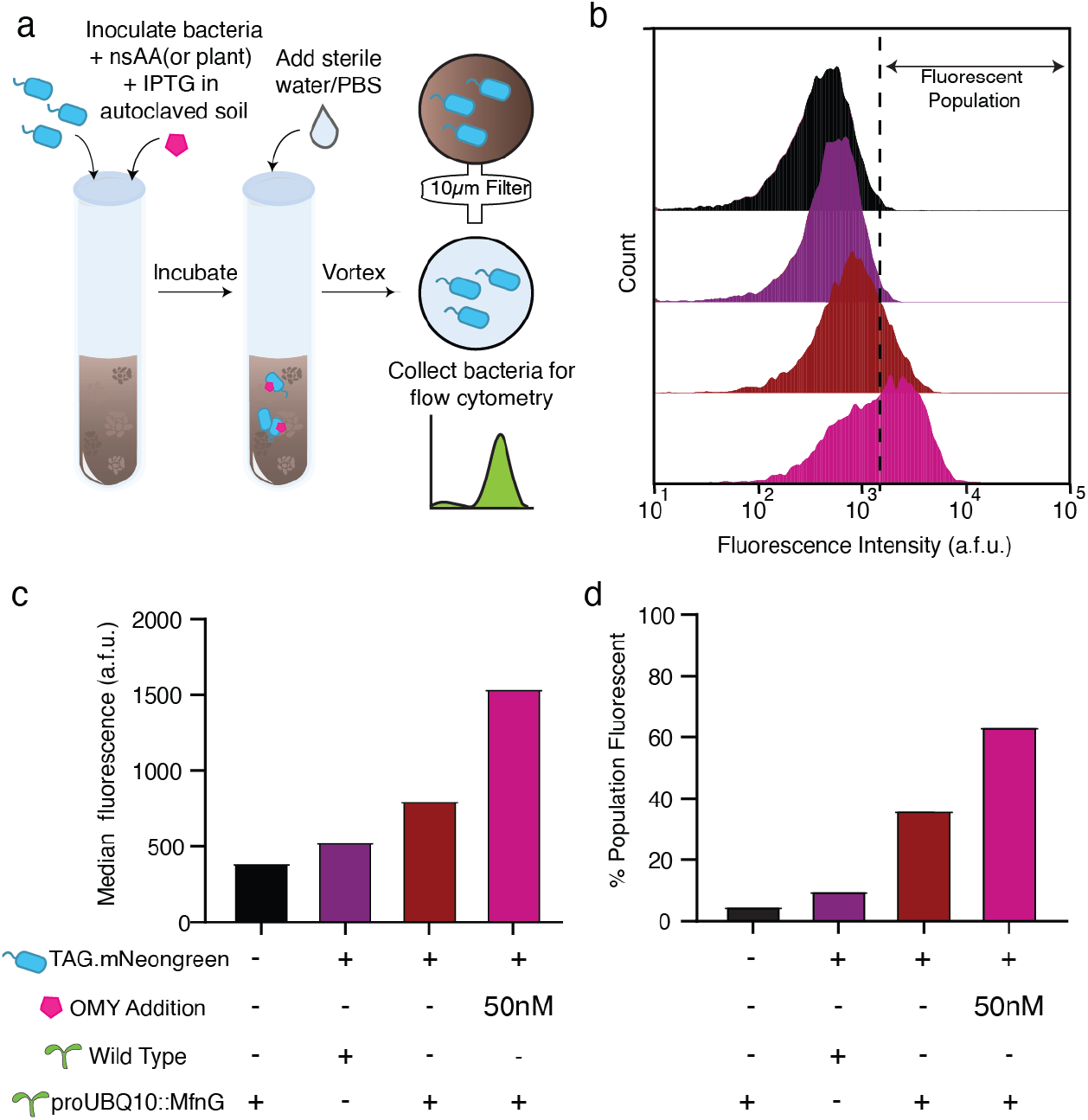
OMY-dependent gene expression in *Bacillus subtilis* UD1022 in soil. **(a)** Workflow for testing OMY-dependent gene expression in soil. Cells were washed, diluted, and inoculated into autoclaved soil with OMY and IPTG as indicated. Bacteria were extracted after 1 or 3 days by adding PBS to soil and filtering through a 10 **μ**m membrane before analysis by flow cytometry. **(b)** Representative flow cytometry histograms containing 10,000 events of UD1022 expressing an OMY-dependent mNeonGreen reporter (TAG.mNeonGreen) extracted from soil planted with proUBQ10::sfGFP-MfnG or wild-type Arabidopsis seedlings. Dashed line indicates the fluorescence of the top 5% of UD1022 wild-type extracted from soil with proUBQ10::sfGFP-MfnG, shown as a reference threshold. Colors correspond to samples as indicated in (c-d). **(c-d)** Median fluorescence (c) and percent of population fluorescent (d) for the same samples as (b).

**Extended Data Fig. 7.**
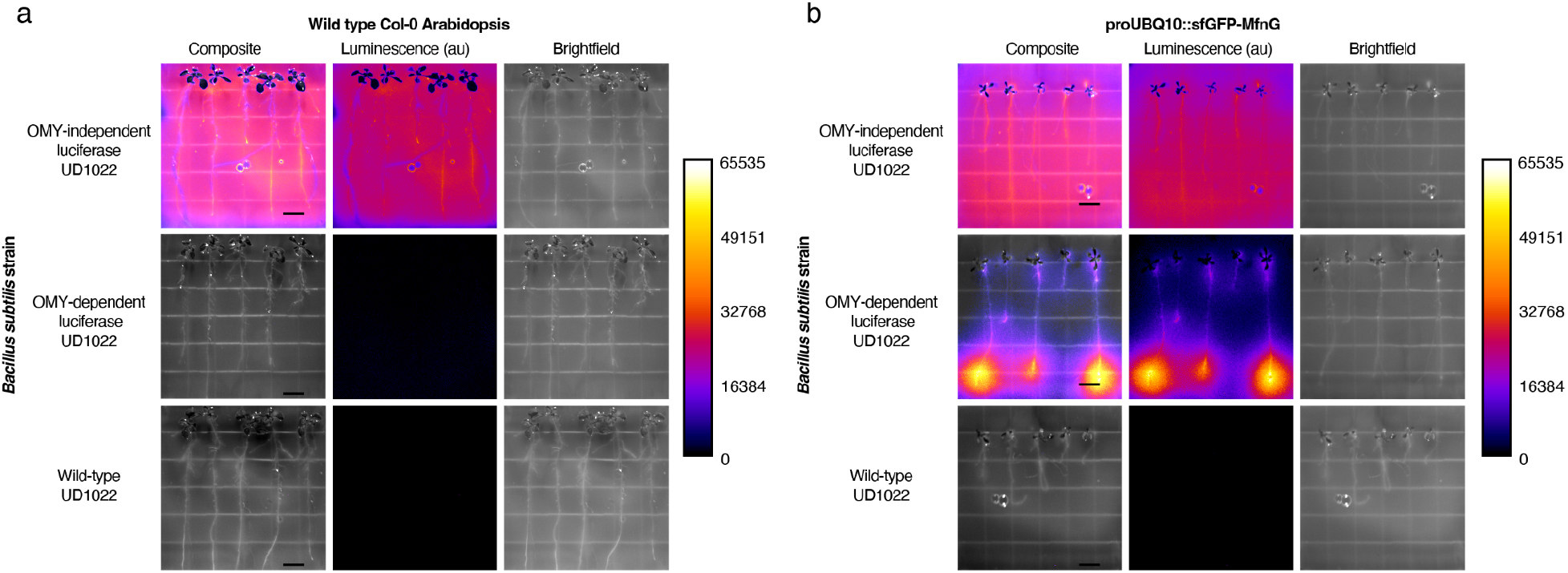
Additional controls for OMY-dependent luciferase expression in *B. subtilis* UD1022 co-cultivated with Arabidopsis. iBright images of *B. subtilis* UD1022 co-cultivated with wild-type Col-0 (a) or proUBQ10::sfGFP-MfnG (b) Arabidopsis seedlings. Composite images overlay luminescence (ChemiBlot channel, 10s exposure) with brightfield (Membrane channel, auto-exposure). Colorbar represents luminescence intensity (Fire LUT). Scale bars, 1 cm.

**Extended Data Fig. 8.**
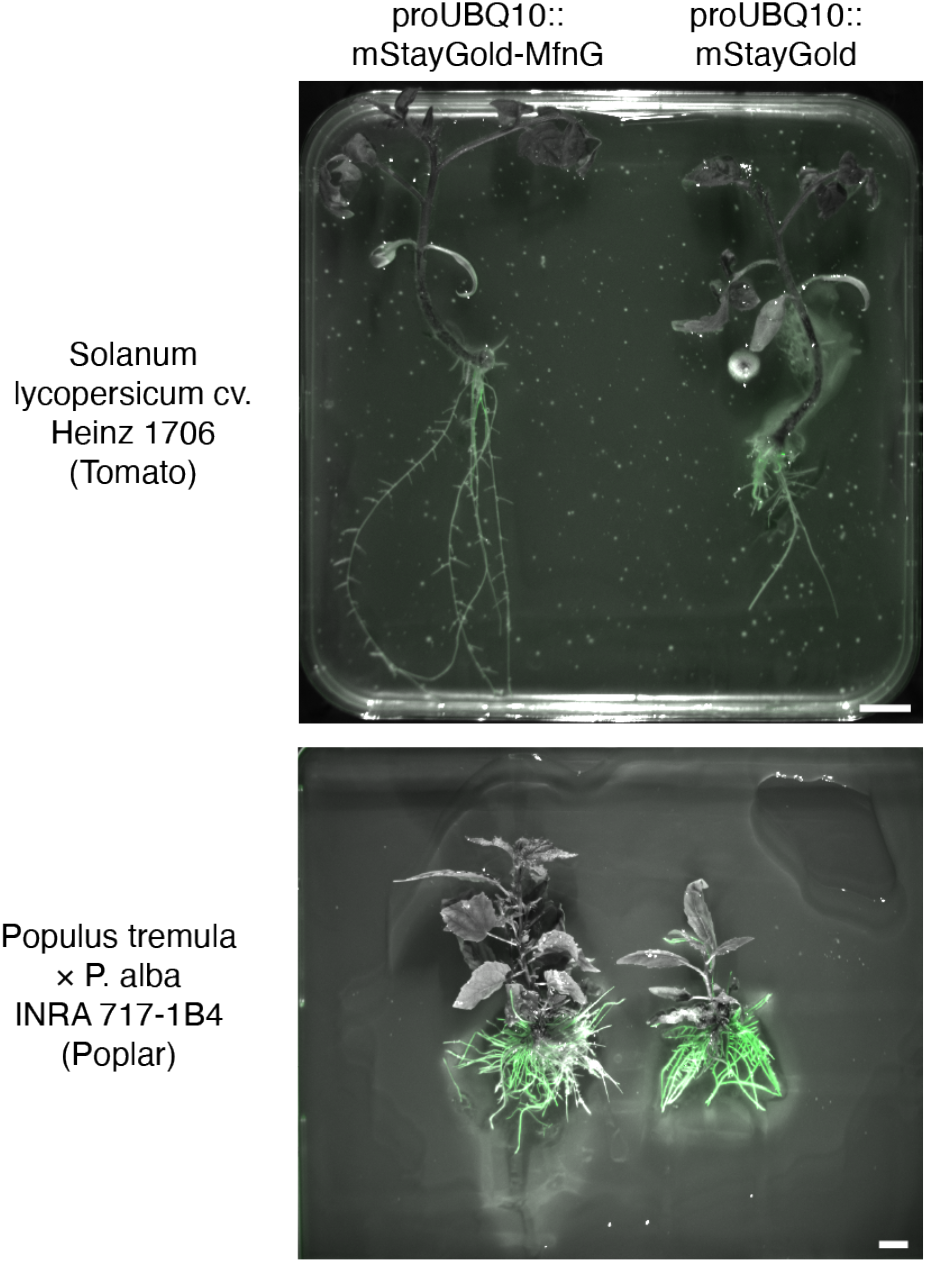
Fluorescence marker confirms successful hairy root transformation. Fluorescence images of tomato (top) and poplar (bottom) composite plants expressing mStayGold in transgenic hairy roots, confirming successful R. rhizogenes-mediated transformation. Images acquired with GFP channel settings in iBright (Ex: 455-485nm, Em: 508-557nm, 15ms exposure). Scale bars, 1 cm.

## Methods

### Bacterial strains and growth conditions

*E. coli* strains NEB10β (Δ(ara-leu) 7697 araD139 fhuA ΔlacX74 galK16 galE15 e14φ80dlacZΔM15 recA1 relA1 endA1 nupG rpsL (StrR) rph spoT1 Δ(mrr-hsdRMS-mcrBC)) and DH5a (F-φ80dlacZΔM15 Δ(lacZY-argF)U169 recA1 endA1 hsdR17(r_K_^−^, m_K_^+^) phoA *sup*E44 λ^−^*thi*-1 *gyr*A96 *rel*A1) were used to construct all plasmids. *Agrobacterium tumefaciens* strain GV3101 containing helper plasmid pSoup (TetR) was used for all Arabidopsis and Nicotiana transformations^78^. *Rhizobium rhizogenes* strain K599 (Intact Genomics 1087) was used for all hairy root transformations. All *B. subtilis* strains were derived from the prototrophic laboratory strain PY79^32,79^ or root isolate UD1022^41^. E. coli and Agrobacterium strains were grown in LB Lenox medium (Sigma L3022) in incubating shakers at 250 r.p.m. (New Brunswick Scientific Innova 44) at 37 °C (E. coli) or 28 °C (Agrobacterium). Antibiotics were used at the following concentrations unless otherwise noted: carbenicillin 100 μg/ml (GoldBio C-103), kanamycin 50 μg/ml (GoldBio K-120), spectinomycin 100 μg/ml (Sigma S4014), gentamycin 50 μg/ml (GoldBio G-400), rifampicin 15 μg/ml (GoldBio R-120), and tetracycline 10 μg/ml (GoldBio T-101). All strains used in this study can be found in **Supplementary Table 1**.

### Recombinant DNA construction

All constructs in this study were generated using Assembly^80^, Golden Gate^81^, and/or overlap extension PCR. DNA parts (promoters, CDS, terminators) were cloned using Gibson assembly into starting plasmids (Level 0), with flanking BsaI cut sites. Cassettes composed of promoters, CDS, and terminators were assembled using Golden Gate reactions to generate constructs (20 µL total reaction volume, 2 µL 10x T4 ligase buffer, 2 µL BsaI-HF, 1 µL T4 DNA ligase, NEB). All constructs and construction primers used in this study can be found in **Supplementary Tables 2-3.** Linearized DNA fragments for cloning in *B. subtilis* were generated with homemade Gibson assembly mixtures combining three distinct pieces: (1) an upstream region containing homologous DNA with the chosen genomic insertion site, an antibiotic resistance cassette, regulator and promoter (if applicable), (2) the reporter of interest, and (3) a downstream region containing homologous DNA with the chosen genomic insertion site including the native terminator.

### *B. subtilis* strain construction

Transformation of genomic, plasmid, or linearized DNA into *B. subtilis* via a natural competency protocol. The parent strains were streaked onto LB (Lennox) plates the day before transformation. Freshly grown colonies were inoculated into 1 mL of transformation media referred to as MC media (100 mM potassium phosphate (monobasic: Sigma-Aldrich P9791, dibasic: Sigma-Aldrich P9666), 3 mM trisodium citrate (Alfa Aesar A12274), 2% D-glucose (TCI America G004825G), 22 mg/mL ferric ammonium citrate (Alfa Aesar A11199), 0.1% casein hydrolysate (Sigma Aldrich 22090), 0.2% L-glutamic acid monopotassium salt (Alfa Aesar A17232)) supplemented with 3 μL of 1 M magnesium sulfate (Sigma Aldrich M7506) and grown for 4 hours at 37 °C in a roller drum (Thermo Scientific Cel-Gro Tissue Culture Rotator Model 1640). Then, 200 μL of culture was mixed with either 1000 ng of linear DNA, 250 ng of plasmid DNA, or 5 μL of *B. subtilis* genomic DNA for transformation. Cultures were incubated at 37 °C in a roller drum for two more hours and then plated on LB agar plates containing the appropriate antibiotics (10 μg/mL kanamycin (Sigma-Aldrich K4000), or 5 μg/mL chloramphenicol (Sigma-Aldrich C0378)). Single colonies were picked and verified by sequencing of purified colony PCR products. For colony PCR, individual *B. subtilis* colonies were suspended in 50 μL Tris-EDTA buffer (10 mM Tris (Sigma-Aldrich 252859), 1 mM EDTA (Sigma-Aldrich E9884), pH adjusted to 8.0 with NaOH/HCl), then frozen at -20°C for 15 min to lyse the cells. 1-2 μL of the thawed cellular mixture was used as a PCR template.

Antibiotic resistance genes were removed according to protocols based on Cre recombinase activity^82^. Briefly, strains were transformed with the pDR244 plasmid. Transformants were streaked on LB agar containing no antibiotics and grown overnight at 42 °C, after which strains were streaked out on media containing either the original antibiotic the strain was resistant to, spectinomycin (final concentration: 100 μg/mL), and no antibiotic to confirm proper curing of the plasmid and antibiotic resistance gene.

### Genomic DNA extraction

*B. subtilis* genomic DNA was extracted using a modified protocol for the Qiagen DNeasy Blood and Tissue Extraction Kit (Qiagen 69504). The strain of interest was streaked out on an LB agar plate and grown overnight at 37°C. A culture was inoculated in LB from a colony and grown to an OD600 of 1.0 at 37 °C shaking for 4-6 h. Cells were then pelleted and resuspended in 180 mL of lysozyme buffer (20 mM Tris-HCl pH 8.0, 2 mM Na EDTA, 1.2% Triton-X with 40 mg/mL of lysozyme added the day of use) and incubated at 37 °C, shaking for 30 minutes. Following lysis, 25 mL of proteinase K and 200 mL Buffer AL from the Qiagen kit were added to the lysate. After a 30 min incubation at 70 °C with shaking, 200 mL of 100% ethanol was added to the mixture. Contents were then added to a spin column and spun down at 20,000 xg for 1 minute. Flow-through was removed and 500 mL of Buffer AW1 was added to the spin column. The column was spun once again at 20,000 xg for 1 min. Flow through was removed, and 500 mL of Buffer AW2 was added. The column was then centrifuged at 20,000 xg for 3 min to remove all buffer. The spin column was then moved to an Eppendorf tube, and 200 mL of Buffer AE added to the spin column. The mixture was incubated at room temperature for 1-5 min, then spun down to elute genomic DNA.

### Measuring ncAA-dependent reporter gene expression in *B. subtilis*

To measure ncAA-dependent reporter gene expression, fresh colonies of engineered *B. subtilis* were picked from LB plates with antibiotics (10 μg/mL kanamycin and 5 μg/mL chloramphenicol) and inoculated into 1 mL of liquid LB media in a 15 mL culture tube containing the same antibiotics. These cultures were grown at 25°C to OD600 0.2 - 0.4 in a roller drum and then sub-cultured depending on the reporter used: (1) For the nanoluciferase reporter strains (strains bMJ252-254, and bMJ257-259), cultures were diluted to an OD ∼0.002 in LB with chloramphenicol, kanamycin and 100 μM OMY (Fisher Scientific M22761G) or (2) For the fluorescent reporter strains (strains bMJ024 and bMJ182), cultures were diluted to an OD of 0.02 in a minimal media with 1/2X Murashige & Skoog Basal Salt Mixture (PhytoTech Labs M524), 1.5% sucrose (Sigma Aldrich S0389), and 10 mM L-glutamate (Acros Organics 156211000) with OMY supplemented at the concentration specified. We leveraged leaky pHyperspank activity for constitutive expression without IPTG induction. Cultures were grown overnight in 300 μL in a 1.3 mL deep well plate (Fisher Scientific 12566611) at 1000 rpm in a shaker with a 2.5 mm orbit (Thermo Scientific 88882005) and incubated at 25 °C unless otherwise specified.

For fluorescent endpoint measurements, 100 μL of the cultures were diluted 1:1 with 100 μL of 1x Phosphate Buffered Saline (1x PBS, Sigma-Aldrich P4417), and then, the absorbance (600nm) and fluorescence (ex: 485nm, em: 525) were read using a Biotek H1 Synergy spectrophotometric plate reader in a black-walled 96-well plate (Fisher Scientific, Greiner-Bio: 07-000-166). Endpoint measurements were also performed in a separate experiment using flow cytometry for Extended Data Fig 6a with settings specified in the flow cytometry methods.

For the nanoluciferase endpoint measurements, cultures were diluted 1:10 with 1x PBS, and OD600 values were measured in a black-walled 96-well plate prior to adding the luciferin reagent. To measure nanoluciferase activity, the Nano-Glo luciferase reagent (Promega N2011) was diluted 1:400 in a 5% solution of the reagent buffer in 1x PBS. The diluted cultures were then mixed in a 1:1 ratio with assay substrate to a total volume of 50 μL in a half-area white-walled 96-well plate (Corning 3885) and read over the course of 10 minutes on the Biotek H1 Synergy plate reader using an integration time of 1 sec, full light emission, top optics, normal read speed with a 100 msec delay and extended dynamic range. The gain was set to 120 after being scaled automatically to the well with the highest luminescence value (the well containing the strain with the wildtype reporter).

For fluorescence measurements by flow cytometry, fluorescence was quantified using a NovoCyte Flow Cytometer. Samples were run with a flow rate of 14 μL min^−1^. *B. subtilis* cells were identified and counted by their clustering in the FSC-SSC space and then gated for singlets. Samples were run until 10,000 events were captured within these gates (**Extended Data Fig. 4**). Single-cell mNeongreen fluorescence levels were measured using a blue laser (488 nM) and the FITC-H filter. The arithmetic median of fluorescence distribution was calculated for each replicate.

### Surfactin Quantification

Strains were grown from LB agar plates to an OD600 of approximately 1.0 in 1/2X MS media supplemented with 1.5% sucrose and 10 mM glutamate containing antibiotics (final concentration of chloramphenicol 5 µg/mL). Strains were then inoculated into 300 µL of the same media, supplemented with OMY as specified, at an initial OD600 of 0.01 in deep well 96-well plates. Samples were grown for 24 hours in an incubator at 37°C and shaken at 1000 rpm before centrifugation to pellet cells. Supernatant was collected for surfactin production quantification.

Surfactin was quantified via high performance liquid chromatography with an Agilent 1100 Infinity model with a Zorbax Eclipse Plus-C18 column with a guard column installed. To isolate surfactin peaks, an isocratic method of acetonitrile with 1% trifluoroacetic acid was used and samples were compared to a commercial standard (Smolecule 24730-31-2). 50 μL of sample was injected and run for 10 minutes, and absorbance was analyzed at 205 nm. Quantification was accomplished by comparing peak areas of three major peaks in the commercial standard.

### Growth kinetics of *B. subtilis*

PY79 expressing an orthogonal aaRS and tRNA (bDS176) was streaked out on LB agar plates with 10 μg/ml chloramphenicol from glycerol stocks. Single colonies were picked and grown in LB media with 10 μg/ml chloramphenicol for about 4 hours at 30°C until they reached an OD600 of approximately 1.0. Cells were then inoculated at an OD600 of 0.01 into 1/2x MS media with 1.5% sucrose and 10 mM L-glutamate with varying concentrations of OMY and 10 μg/ml chloramphenicol. Growth was monitored using OD600 measurements on a Biotek Synergy H1 plate reader for 24 hrs with continuous shaking at 30°C.

### Flow cytometry of microbes extracted from soil

Soil (Espoma Organic Potting Mix 00050197701044) samples were prepared by weighing out 1 g of soil into glass culture tubes (18 x 150 mm) and autoclave sterilizing for 20 minutes at 121°C. Bacteria were grown by picking fresh colonies from LB agar into liquid LB media, both containing antibiotics (chloramphenicol (final concentration: 5 µg/mL) and kanamycin (final concentration: 10 µg/mL)). These cultures were grown at 37°C to OD_600_ of 0.2–0.4 in a roller drum. After, cells were washed 6 times in phosphate-buffered saline (PBS) before diluting to a final OD_600_ of 0.5 and 1mM IPTG and 500 µM of OMY was added. One mL of cell suspension was mixed into each soil sample. Samples were incubated at 25°C in the dark with a damp paper towel to prevent samples from drying out. After 1-3 days, cells were harvested by adding 5mL of sterile PBS to the bacteria-soil mixture, vortexing, and filtering through a 10 µm syringe filter. Fluorescence was quantified using a NovoCyte Flow Cytometer. Samples were run with a flow rate of 14 μL min^−1^. *B. subtilis* cells were identified and counted by their clustering in the FSC-SSC space; gates for singlets were drawn on PY79 and UD1022 samples grown in liquid LB and then applied to samples extracted from soil (**Supplementary Data Fig. 2**). Samples were run until 10,000 events were captured within these gates. Single-cell mNeongreen fluorescence levels were measured using a blue laser (488 nM) and the FITC-H filter. The arithmetic median of fluorescence distribution was calculated for each replicate.

### Co-cultivation of microbes and plants in soil

Soil was sieved through a mesh strainer and 0.5 g was weighed out into a 15 mL conical tube. Soil and tubes were autoclaved at 121 °C for 20 min. Pre-sterilized Arabidopsis seeds were imbibed in 1.5 mL sterile water and cold-stratified in the dark for 3 days at 4 °C. After stratification, seeds were sown into autoclaved soil and incubated at 22 °C in 60% relative humidity with a 16/8 h photoperiod for plants to germinate and grow. *B. subtilis* were prepared for co-cultivation by harvesting exponential phase cultures grown in LB and washing six times in sterile water. After washing, cultures were diluted to 0.1 OD600 and mixed with 1 mM IPTG. 500 μL of the cell solution was added to the soil containing 11 day old seedlings. Plants and bacteria were co-cultured for 3 days before samples were harvested for flow cytometry analysis as described above.

### Plant growth conditions

*Nicotiana benthamiana* plants were grown and maintained in a Percival growth chamber (AR100L3) at 24 °C in 12h/12h light/dark cycles with 70% relative humidity. Light was provided by fluorescent white lights at a photon flux density of ∼150-200 μmol m⁻² s⁻¹. Plants were grown in 4-inch pots with soil containing mycorrhizae, peat moss, and perlite (PRO-MIX BX 10381RG).

*Arabidopsis thaliana* plants were grown and maintained in a walk-in growth chamber (Percival AR-712L4) at 22/18 °C in 16/8-h light/dark cycles with 60% relative humidity for transformation and seed propagation. Light was provided by LED arrays at a photon flux density of ∼145 μmol m⁻² s⁻¹. Plants were grown in 4-inch pots with soil containing mycorrhizae, peat moss, and perlite (PRO-MIX BX 10381RG).

*Arabidopsis thaliana* plants for imaging were grown and maintained in a separate growth chamber (Percival CU36L4) at 22/18 °C in 16/8-h light/dark cycles. Light was provided by white fluorescent lights at a photon flux density of ∼145 μmol m⁻² s⁻¹. Plants for imaging and root structure analysis were plated as seeds onto sterile MS media containing 0.6% Gelzan (Bioworld 30634003), 1X Murashige & Skoog Basal Salt Mixture (PhytoTech Labs M524), 1% sucrose (Sigma S0389), 0.05% MES (Sigma-Aldrich M8250), pH 5.7. Prior to plating, seeds were surface-sterilized with 70% ethanol for 5 min, followed by 20% bleach and 0.1% (v/v) Tween-20 (Sigma-Aldrich P7949) for 5 min, then washed three times with water for 2 min each. Seeds were cold-stratified for 3 days at 4 °C in the dark, either on plate or in water prior to plating, before transfer to the growth chamber. Plates were sealed with micropore tape (3M 1530-0) to allow for gas exchange.

### Transient expression of constructs in *Nicotiana benthamiana* leaves

*Agrobacterium tumefaciens* GV3101 cultures harboring the desired plasmids were inoculated from frozen glycerol stock (18% glycerol) into 14 ml culture tubes (VWR 47729-566) containing 2 mL of LB media with antibiotics. Cultures were grown for 24 h at 28°C and 250 r.p.m. The next day, cultures were back diluted to an OD600 of 0.02 in fresh LB media with antibiotics and grown for another 24 h at 28°C and 250 r.p.m. Cells were then collected by centrifugation (8 min, 3000g), resuspended in 1 mL infiltration buffer (10 mM MgCl_2_ (Sigma M8266), 10 mM MES pH 5.7 (Sigma M8250), 150 μM acetosyringone (Cayman 23224)), and incubated at room temperature for 3 h. Agrobacterium strains were then mixed to generate samples for leaf infiltration. The strain containing the sfGFP-MfnG construct was added to the mixture at an OD600 of 0.3, and a strain containing the p19 suppressor of silencing was added at an OD600 of 0.2. Final sample volumes were adjusted to 1 mL using infiltration buffer. Samples were infiltrated into the abaxial side of the N. benthamiana leaves using a 1 mL needleless syringe (Fisher Scientific 22-253-260). Infiltrated plants were returned to normal growth conditions for ∼72h. Leaf discs were then harvested from infiltrated and uninfiltrated areas of the leaf using the back of a pipette tip (P20 Rainin 17005873) for metabolite extraction and LC-MS analysis.

### Generation of *Arabidopsis* transgenic lines

Heritable transgenic lines of *Arabidopsis* were generated by *Agrobacterium tumefaciens*-mediated floral dip of *Arabidopsis thaliana* Columbia (Col-0) ecotype^83^. Briefly, *A. tumefaciens* cultures were inoculated from frozen glycerol stock (18% glycerol) into 14 ml culture tubes containing 2 mL of LB media with antibiotics. Cultures were grown for 24 h at 28°C and 250 r.p.m. After incubation, 250 μL of bacterial culture was plated on 15 cm round LB agar plates containing antibiotics and incubated for two nights at 28°C. Next, the cells were scraped from plates and resuspended in 200 mL transformation buffer (50 g/L sucrose, 250 μL/L Silwet L-77 (Bioworld 30630216)). Arabidopsis inflorescences were briefly submerged in the bacterial solution and then transferred to a dark, humid container where they were allowed to incubate overnight. The following day, transformed plants were returned to normal growth conditions where they continued to grow until they produced seeds. Transgenic seeds were screened using fluorescence microscopy. All constructs used for *Arabidopsis* transformation contain FastRed, a seed-specific TagRFP marker, which enables selection of T1 transgenic plants by screening for red fluorescence in dry seeds^84^. For each construct, we screened multiple independent T1 events and selected one or more representative lines, denoted by event number, for further characterization.

#### Generation of OMY-producing tomato hairy roots

Tomato seeds (*Solanum lycopersicum* cv. Heinz 1706) were surface-sterilized by briefly soaking (<1 min) in 70% (v/v) ethanol, followed by incubation for 15 min in a sterilization solution containing 2.5% (v/v) sodium hypochlorite (prepared by diluting 3 mL of 8.25% commercial bleach in 10 mL distilled water) and 0.05% (v/v) Tween-20. Seeds were then washed five times with sterile distilled water and drained thoroughly. Sterilized seeds were soaked overnight in sterile distilled water at room temperature and subsequently placed on sterile MS media plates containing 0.6% Gelzan, 0.5X Murashige & Skoog Basal Salt Mixture and 0.05% MES at pH 5.7. Seedlings were grown in a growth chamber (Percival CU36L4) in standard conditions (22/18 °C, 16/8-h light/dark cycles) with a photon flux density of approximately 145 μmol m⁻² s⁻¹ for 8 days. The radicle and lower hypocotyl of 8-day-old seedlings were excised, retaining the apical 4 cm of the hypocotyl. The freshly cut wound surface was inoculated by coating with transgenic *Rhizobium rhizogenes* strain K599 harboring the construct of interest. The bacteria were pre-cultured on YEB plates supplemented with 100 µg/mL kanamycin and 200 µM acetosyringone, then applied to the wound surface using a sterile pipette tip. Inoculated seedlings were transferred to fresh MS gelzan plates and co-cultivated for 3 days under the same growth chamber conditions. After co-cultivation, seedlings were transferred to MS gelzan plates supplemented with 400 mg/L timentin (Goldbio T-104) to eliminate *R. rhizogenes*. Seedlings were subcultured onto fresh timentin-containing MS plates every 14 days. Adventitious roots emerging initially from the wound site are not transgenic; transgenic hairy roots were first observed approximately 2 weeks after inoculation. Plants were maintained for 6 weeks post-inoculation to allow sufficient transgenic root biomass to accumulate before use in downstream luminescence assays.

#### Generation of OMY-producing poplar hairy roots

*Populus tremula × P. alba* (clone INRA 717-1B4) plants were grown under aseptic conditions in MS agar medium (0.5X Murashige & Skoog Basal Medium with Vitamins (PhytoTech Labs M519), 4 g/L sucrose, 7 g/L phytoagar (Bioworld 40100072), 0.05% MES, pH 5.8) in a growth chamber (Percival CU36L4) under standard conditions (22/18 °C, 16/8-h light/dark cycles with a photon flux density of approximately 145 μmol m⁻² s⁻¹photoperiod). Apical explants were prepared for transformation by excising plant segments with two to three leaves using a sterile scalpel. The freshly cut wound surface was inoculated by dragging it across a lawn of transgenic *Rhizobium rhizogenes* strain K599 harboring the construct of interest. The bacteria were pre-cultured in 5 mL liquid LB medium supplemented with appropriate selection antibiotics at 28°C for 24 h, then plated onto solid LB medium containing antibiotics and 200 uM acetosyringone. Plates were incubated at 28°C for 48 h to establish a bacterial lawn for inoculation. Inoculated explants were immediately inserted into antibiotic-free MS medium and co-cultivated for 3 days under the same growth chamber conditions as before. Following co-cultivation, explants were transferred to fresh MS medium supplemented with 200 mg/L timentin and 200 mg/L carbenicillin to eliminate *R. rhizogenes*. Transgenic hairy roots were first observed approximately 4 weeks post-inoculation. At 5 weeks post-inoculation, transformed plants were excised from the selective medium for co-cultivation with *B. subtilis* luciferase reporter strains for luminescence assays.

### Fluorescence imaging of *Arabidopsis* transgenic lines

Fluorescence images of *Arabidopsis* lines expressing sfGFP-MfnG constructs were collected using a Leica THUNDER widefield fluorescence microscope (M205 FCA) with the ET GFP filter (Leica 10447408, Ex470/40, Em525/50) at 150ms or 500ms exposure as noted. All image processing was performed using the Fiji software. Processing steps include: setting thresholds for minimum and maximum brightness, applying LUTs (Green for green fluorescence, Fire for luminescence), generating colorbars, and merging channels.

#### Root exudate collection and metabolite extraction

Root exudate from wild-type Col-0 and transgenic MfnG-expressing *Arabidopsis* was collected in MS media for metabolite quantification by LC-MS. Seeds were sown on sterile MS plates (0.5X MS basal salts, 1% sucrose, 0.6% Gelzan, pH 5.7), stratified for 3 days in the dark at 4°C, then moved to the growth chamber (Percival CU36L4) and grown under standard conditions (22/18 °C, 16/8-h light/dark cycles). After 6 days, seedlings were transferred to 3’ x 3’ x 4’ Magenta boxes (Bioworld 30930007 GA-7) for hydroponic growth in 12 mL of sterile liquid media (0.5X MS basal salts, 1% sucrose, pH 5.7). Magenta boxes were sealed with micropore tape and incubated for 48 hours under standard *Arabidopsis* growth conditions (see Plant growth conditions) on an orbital shaker at 50 rpm. After 48 hours, 10 mL of media was collected from each box with a serological pipette, syringe-filtered through a 0.45 µM filter, and transferred to 15 mL centrifuge tubes. To prepare samples for lyophilization, tubes were frozen at -80°C for 30 min, then moved to dry ice. Tube caps were replaced by Kimwipes secured over the opening with rubber bands for lyophilization in a Labconco lyophilizer (Labconco 710612010) for 24 hours. Samples were stored at -80°C after lyophilization until metabolite extraction. To extract metabolites, 500 µL of 80% methanol (v/v) was added to each sample, briefly vortexed, sonicated (Branson 2800) for 15 min at the High setting, and centrifuged for 10 min (ThermoScientific Sorvall X4R Pro TX1000 rotor) at 4000 rpm. For each sample, 400 µL of supernatant was sampled for LC-MS analysis.

#### LC-MS analysis to quantify OMY production

LC–MS samples were analyzed on an Agilent 1260 HPLC system coupled to an Agilent 6520 Q-TOF mass spectrometer. Separation was done using a Gemini 5-μm NX-C18 110-Å column (2 × 100 mm; Phenomenex) with a mixture of 0.1% formic acid in water (A) and 0.1% formic acid in acetonitrile (B) at a constant flow rate of 400 μl per min at room temperature. The injection volume was 3 μl for experimental samples and 1 μl for metabolite standards. The following gradient of solvent B was used: 1% 0–1 min, 1%–8% 1–3 min, 8%–97% 3–15 min, 97-1% 15– 16 min, 1% 16-22 min. MS data were collected using electrospray ionization (ESI) in positive mode with a mass range of 50–1,200 m/z and a rate of one spectrum per second. The ionization source was set as follows: 325°C gas temperature, 10 l min^−1^ drying gas, 35 psi nebulizer, 3,500 V VCap, 150 V fragmentor, 65 V skimmer, and 750 V octupole 1 RF Vpp.

Data were collected with Agilent MassHunter Workstation Data Acquisition and analysed by MassHunter Qualitative Analysis 10.0. Extracted ion chromatograms (EICs) were extracted for [M+H]^+^ ions using 196.067 m/z for OMY and 182.0800 m/z for tyrosine, with an error of 20 ppm. Counts per minute data were exported to CSV files for further data analysis and visualization. For conversion of integrated peak values to OMY concentration, an OMY calibration curve was run at the same time as the samples with concentrations 0.5 µM, 1 µM, 2 µM, 5 µM, 25 µM, and 100 µM with two technical replicates per concentration. OMY stock for LC-MS was prepared at 10 mM with MilliQ water without adjusting pH.

#### Root phenotyping

For analysis of root length, lateral root density, and gravitropism, *Arabidopsis* seeds were germinated and grown on MS media containing 0.6% Gelzan (Bioworld 30634003), 1X Murashige & Skoog Basal Salt Mixture (PhytoTech Labs M524), 1% sucrose (Sigma S0389), 0.05% MES (Sigma-Aldrich M8250), pH 5.7. For exogenous treatment of seedlings with O-methyl-L-tyrosine (OMY), appropriate volumes of filter-sterilized 500 mM stock solution of OMY (Chem-Impex 06251) was mixed into the MS gelzan media prior to gelling. OMY stock solution was prepared in MilliQ water with 10 M KOH added until all OMY was dissolved (final pH ∼11). For exogenous treatment of seedlings with standard amino acids, filter-sterilized L-tyrosine (1 mM in MilliQ, pH adjusted to 9.7 with 10M KOH, overnight stirring, Research Products International T68500) and L-phenylalanine (11 mM in MilliQ, Research Products International P20260) stocks were added to MS gelzan media prior to gelling. Seeds were sown and germinated directly on OMY and amino acid treatment plates. After growth, plates were imaged using a dual light flatbed scanner (Epson V800, 800 dpi) at different timepoints post-stratification provided in results and figure captions. Total root length was quantified using the Fiji SmartRoot tool^85^ and measured from the hypocotyl-root junction to the root tip. Plants that did not germinate were discounted from the analysis. The number of emerged lateral roots was counted from the scanned images. Lateral root density calculated as the number of emerged lateral roots divided by primary root length. Gravitropism score is calculated as the vector length of the root on the plate divided by primary root length.

#### Co-cultivation of plants and bacteria on plates for luciferase imaging of OMY incorporation

To create the bacterial-agar matrix for co-cultivation of plants and bacteria, *B. subtilis* strains were streaked from frozen glycerol stocks onto LB agar plates with appropriate antibiotics and incubated at 37 °C overnight. Colonies were inoculated into 50 mL LB without antibiotics in 200 mL flasks and incubated at 37 °C with 250 rpm shaking. After cultures reached OD600 ∼1 (3-4 h), they were decanted into 50 mL centrifuge tubes (Corning 352070) and washed three times with 25 mL sterile 1X PBS (Sigma-Aldrich P4417-100TAB), each time pelleting at 4000 rpm for 10 min (ThermoScientific Sorvall X4R Pro TX1000 rotor). The final pellet was resuspended in 1 mL 1X PBS. MS agar media (0.5X MS basal salts, 1% sucrose, 1% agar, 0.05% MES, pH 5.7) was prepared, autoclaved, and allowed to cool to ∼55 °C. Bacteria were then mixed into the media at a final OD600 of 0.1. When appropriate, β-estradiol was mixed into the MS agar at this stage (1:1000 v/v) to a final concentration of 20 μM. After mixing, the bacteria-agar mixture was immediately pipetted into sterile petri dishes. For Arabidopsis seedlings, 30 mL of mixture was added to 100 mm square plates (Genesee 26-275); for tomato composite hairy roots, 50 mL to 120 mm square plates (Greiner Bio-One 688161); for poplar composite hairy roots, 250 mL to 245 mm square plates (Corning 431111).

Once the bacteria-agar matrix cooled, plants were transferred onto the plates for co-cultivation using sterile forceps. Plant roots were placed directly onto the surface of the solidified bacteria-agar matrix, allowing root-bacteria contact to occur at the agar surface. Arabidopsis plants or composite plants were placed onto each plate, which were then sealed with micropore tape, wrapped in foil from the bottom up to the top of the shoot, and returned to the growth chamber for co-cultivation in standard conditions. A schematic of the co-cultivation setup is provided in

#### Supplementary Fig. 1

After 4 days of co-cultivation (5 days for poplar), luciferin was applied to the plate as an agarose film. This luciferin-agarose mixture is overlaid on top of the co-cultivated roots and bacteria to deliver the luciferase substrate for imaging. To create this film, 0.8% agarose (Research Products International, A20090) was dissolved in water and cooled to ∼55 °C. NanoFuel (NanoLight Technology 324-50, Arabidopsis assays) or NanoGlo (Promega N1110, tomato and poplar assays) substrates were diluted 1:25 into their respective buffers, then added to the molten agarose solution for a final dilution of 1:500. Luciferin-agarose mixtures were immediately applied to co-cultivation plates by slowly pipetting a thin surface layer over plants roots and bacteria (10 mL for Arabidopsis assays, 15 mL for tomato, 60 mL for poplar). Co-cultivation plates were immediately imaged with an iBright Imaging System (Invitrogen FL1500) with the following settings: brightfield (Membrane channel, auto-exposure), luminescence (ChemiBlot channel, 30s exposure unless otherwise indicated), fluorescence (RFP Ex: 515-545nm, Em: 568-617nm, 15ms exposure; GFP Ex: 455-485nm, Em: 508-557nm, 15ms exposure). All image processing was performed using the Fiji software. Processing steps include: setting thresholds for maximum brightness, applying LUTs (Green for green fluorescence, Fire for luminescence), generating colorbars, and merging channels.

For β-estradiol induction, 20 mM β-estradiol was mixed into the MS gelzan-bacteria media prior to gelling at 1:1000 v/v. β-Estradiol (Sigma-Aldrich E2758) stock solution was prepared in DMSO (Sigma-Aldrich 34869).

#### Quantification and analysis of luminescence intensity

Raw luminescence (ChemiBlot channel) images (16-bit) were used for quantification with Fiji. Square ROIs were drawn on the corresponding brightfield image (Membrane channel) following grid borders on the petri dish. ROIs were propagated to the luminescence image and the Plot Profile tool was used to plot the vertically averaged pixel intensity at distances along the x-axis, starting with 0 cm at the left extreme.

Further data analysis was implemented in Python 3 using numpy, scipy (v1.11), and pandas. Local maxima (peaks) in the luminescence intensity profile corresponding to the location of individual seedlings were extracted as follows. Each profile was smoothed with a uniform (boxcar) moving-average filter (kernel width = 1/10 of the expected inter-peak spacing) to reduce noise. Candidate peaks were identified using *scipy.signal.find_peaks* with a minimum inter-peak distance set to half the expected spacing. The prominence threshold parameter was iteratively lowered from 30% to 1% of the intensity value range until at least n (the number of seedlings) peaks were found. Only the n most prominent peaks were retained. If fewer than n peaks were detected, the largest inter-peak gap was bisected and the local maximum within that gap was added. Each detected peak was then snapped to the local maximum of the unsmoothed profile within ±3 data points (∼0.08 cm). For PY79 negative-control plates, since no peaks were expected, peak positions were assigned to fixed equidistant intervals along the x-axis.

For each peak at position x₀, two midpoints were defined: x₁ halfway between x₀ and the adjacent peak to the left (or the profile start), and x₂ halfway to the adjacent peak to the right (or the profile end). The peak (maximum) intensity y was taken as the exact (unsmoothed) value at x₀. The baseline intensities y₁ and y₂ were computed as the mean value within ±0.1 cm of x₁ and x₂, respectively, and averaged to yield the base intensity y₀ = (y₁ + y₂)/2. The normalized luminescence ratio (fold change) was calculated as y/y₀. Peak and midpoint locations are annotated in **Supplementary Fig. 3**. Claude Science Opus 4.6 was used in the development of the Python analysis script.

### Proteomics

All surfaces and equipment were cleaned with ethanol to prevent keratin contamination. Methanol was avoided to prevent artificial methylation.

#### Protein extraction

Arabidopsis seedlings were grown on 1X MS gelzan plates and harvested at 14 days. Shoot and root tissue were separated with a razor blade and placed into separate 2 mL Safe-Lock tubes (Eppendorf 022363344) with forceps. Samples were immediately weighed and flash-frozen in liquid nitrogen, then stored at -80 °C until extraction. Sample collection data is provided in **Supplementary Table 4.** To homogenize tissue, one metal bead was added to each tube and samples were lysed for 2 min at 30 Hz (RETSCH MM 400 Mixer Mill). 2X SDS sample loading buffer (125 mM Tris-HCl, pH 6.8 (VWR 97061-796), 4% sodium dodecyl sulfate (Sigma 1125330250), 20% glycerol (Sigma G5516), 5% v/v beta-mercaptoethanol (Sigma M3148)) was added to tissues at 3:1 v/w. Samples were then boiled for 5 min at 95 °C (Thermolyne DB28125 Dri-Bath-Marshall Scientific). After cooling, samples were spun down in a benchtop centrifuge at 13200 rpm for 20 min, and supernatants were extracted into new tubes for gel purification.

#### Gel electrophoresis and staining/de-staining

50 µL of each supernatant was loaded into the well of a polyacrylamide gel (BioRad 4561094). A buffer lane was left between each sample to prevent cross-contamination. Samples were run in 1X Tris/Glycine/SDS buffer (Bio-Rad 1610732) at 100V until 5-10 mm into the resolving gel. The gel was then stained for 6 h in a methanol-free staining buffer to prevent artificial methylation. 50 mL of buffer was prepared with the following in ddH_2_O: 10.5 g of citric acid monohydrate (final concentration 1M, Millipore Sigma C1909), 2.5 mL glacial acetic acid (final 5%, Fisher chemicals AA36289AP), 5 mL of Stainer B and 10 mL of Stainer A from the Invitrogen™ Colloidal Blue Staining Kit (ThermoFisher Scientific LC6025). Gels were destained with HPLC-grade water overnight.

#### Protein in-gel digestion and peptide purification

Following destaining, whole protein bands were excised from the gel using a scalpel inside a laminar flow hood, limited to a maximum length of 10 mm. Excised bands were diced into small 1 mm^2^ pieces in the presence of a droplet of mass spectrometry-grade water to facilitate cutting, and the pieces were transferred into 0.65 mL low-retention microcentrifuge tubes (Fisher Scientific 13698793). The gel pieces were destained by three wash cycles of adding 400 µL of buffer B1 (25 mM ammonium bicarbonate (Millipore Sigma A6141) in 50% acetonitrile (Fisher Chemicals A998SK-4) v/v), vortexing for 10 min, and removing the supernatant. This was followed by one 5-min wash cycle of buffer A1 (25 mM ammonium bicarbonate in ddH_2_O) and one 15-min wash cycle with buffer B1, until the blue dye largely disappeared from the gel pieces.

For reduction and alkylation, disulfide bonds were reduced by incubating the gel pieces in 100 µL of 10 mM dithiothreitol (Millipore Sigma D0632) (prepared from 500 mM stock in 25 mM ammonium bicarbonate) at 56 °C for 1 hour. After cooling to room temperature, 100 µL of 50 mM iodoacetamide (Millipore Sigma I1149, prepared from 500 mM stock in 25 mM ammonium bicarbonate) was added, and alkylation was allowed to proceed in the dark for 45 minutes at room temperature. The reagents were removed, and the gel pieces were washed three times with 200 µL of buffer B1 with 10-min vortexing steps. Gel pieces were then evacuated to complete dryness with a SpeedVac system (Speedvac concentrator: Thermoscientific SavantSPD121P, Refrigerated vapor trap: Thermoscientific Savant RUT5105, Vacuum pump: Thermoscientific OFP400).

The dried gel pieces were rehydrated and digested by adding 10-50 µL of trypsin to just barely cover the gel pieces. Trypsin (Thermofisher Scientific 20233) was freshly diluted to 5 ng/µL in buffer A1 before use. Following a 5-min incubation with trypsin, add additional buffer A1 as needed to ensure the gel pieces remained entirely submerged in solution. Digestion was allowed to proceed overnight at 37 °C.

Peptides were extracted by adding 100 µL of buffer B2 (50% acetonitrile and 0.1% formic acid (Fisher Chemicals A117-50) v/v in HPLC-grade water). The samples were vortexed for 15 minutes, and the peptide-rich supernatant was carefully transferred to fresh low-retention tubes while avoiding gel carryover. This extraction step was performed three times total. The combined supernatants were centrifuged at 20,000 rpm (Eppendorf 5417C) for 15 minutes to pellet any debris, and ∼400 µL of supernatant was transferred into fresh tubes. The sample tubes were flash-frozen in liquid nitrogen for 2 minutes with lids open, then placed into a SpeedVac to dry completely (about 2 h).

Peptides were then desalted with ZipTip OMIX C18 tips (Agilent A57003100). The dry peptide residues were reconstituted in 20 µL of buffer A2 (0.1% formic acid v/v in HPLC-grade water) and centrifuged at 20,000 rpm for 15 minutes. 20 µL of supernatant was transferred to a fresh tube. The C18 matrix was activated by aspirating and discarding 20 µL of buffer B2, then equilibrated by aspirating and discarding 20 µL buffer A2 four times. The reconstituted peptide sample was bound to the resin via eight aspirate-dispense cycles. The matrix was then washed with buffer A2 eight times to remove salts. The purified peptides were eluted directly into fresh low-retention tubes by aspirating and dispensing buffer B2 eight times. Continuous wetting of the ZipTip was maintained during the operation to prevent collapse of the matrix. The eluted samples were dried completely in a SpeedVac (about 30 min) and stored at - 80 °C prior to mass spectrometry analysis.

#### LC-MS/MS analysis

LC-MS/MS was carried out on an Orbitrap Eclipse mass spectrometer (Thermo Fisher), equipped with an Easy LC 1200 UPLC liquid chromatography system (Thermo Fisher). Peptides were first trapped using a trapping column (Acclaim PepMap 100 C18 HPLC, 75 μm particle size, 2 cm bed length), then separated using analytical column AUR3-25075C18, 25CM Aurora Series Emitter Column (25 cm x 75 µm, 1.7 µm C18) (IonOpticks). The flow rate was 300 nL/min. Peptides were eluted by a gradient from 3 to 28 % solvent B (80 % acetonitrile, 0.1 % formic acid) over 106 min and from 28 to 44 % solvent B over 15 min, followed by a short wash (15 min) at 90 % solvent B.

For data acquisition, precursor scan was set to mass-to-charge ratio (m/z) 375 to 1600 (resolution 120,000; AGC 200,000, maximum injection time 50ms, normalized AGC target 50%, RF lens 30%) and the most intense multiply charged precursors were selected for fragmentation (resolution 15,000, AGC 5E4, maximum injection time 22 ms, isolation window 1.4 m/z, normalized AGC target 100%, include charge state=2-8, cycle time 3 s). Peptides were fragmented with higher-energy collision dissociation (HCD) with a normalized collision energy (NCE) of 27. Dynamic exclusion was enabled for 30s.

#### Data search with MSFragger & Perseus

MS/MS spectra were searched separately against the Araport11 database *Arabidopsis thaliana* (https://www.arabidopsis.org/), concatenated with decoy protein sequences using the MSFragger 4.1 software by including O-methylation at tyrosine (delta mass of +14.01565) as a variable modification with default criteria to obtain Label Free Quantitative (LFQ) intensities. The search results were analyzed separately in Perseus (version 2.0.10.0). The processing in Perseus was as follows: MSFragger intensities were log2-transformed. Only proteins that had at least three valid values in at least one group (WT or proUBQ10::sfGFPMfnG) were kept. The remaining missing MaxLFQ intensities were then imputed from a normal distribution that is downshifted by 1.8 and a width of 0.3 column-wise. A two-sample t-test was conducted with a permutation-based (n=250) FDR = 0.05 and S_0_ = 1.5.

#### Graphing and statistical analysis

Graphs were constructed using GraphPad Prism and Python packages. Flow cytometry histograms were constructed in the NovoExpress software. Mann Whitney and Kruskal Wallis (two-tailed, no multiple comparison correction) significance tests were computed in Prism. Relative IC50 curves were fitted in Prism from median values.

## Acknowledgements

We thank collaborators who provided plant seeds and bacterial strains — specifically, Jung-Gun Kim and Mary Beth Mudgett at Stanford University for providing tomato seeds, Chris Dervins and Matias Kirst from the University of Florida for providing poplar cuttings and a propagation protocol, Harsh Bais from the University of Delaware for providing *B. subtilis* UD1022, and Devon Stork for providing *B. subtilis* constructs and genetic code expansion protocols. We thank Cathy Fromen and Emma Sudduth for access to and training on the flow cytometer at the University of Delaware. We thank Natalie Farny at Worcester Polytechnic Institute for providing protocols and guidance on microbial extractions from soil. We thank Yaereen Dho, Jack Liu, Sarah Niehs, Diego Wengier, and Elizabeth Sattely for access to and training on the lyophilizer and LC-MS at Stanford University, as well as guidance on LC-MS experiment design and data interpretation. We thank Brian Caliando for the luciferin agarose protocol.

## Author Contributions

### Contributions

V.Z., M.A.J., A.M.K., and J.A.N.B conceptualized the project and contributed to experimental design, data analysis, and data interpretation. V.Z. generated transgenic Arabidopsis lines, quantified OMY production from plant lines, performed phenotypic characterization, and performed image processing and analysis. V.Z. and A.G. designed and performed plant-microbe co-cultivation assays. A.G. contributed to the phenotypic characterization of transgenic Arabidopsis lines. M.A.J. and A.C. designed bacterial strains containing genetic code expansion technology and performed preliminary characterization of bacterial strains. R.I. designed bacterial strains for hairy root transformation and generated tomato hairy roots. A.A.J. generated poplar hairy root and contributed to co-cultivation assays. A.C. performed incorporation assays using flow cytometry in soil and with the engineered plant. S.S.K. and S.X. performed proteomics analysis to assess proteome-wide incorporation of OMY in Arabidopsis. A.M.F. designed and initially screened constructs for O-methyl-transferase expression. V.Z. and J.A.N.B. wrote the manuscript with contributions from M.A.J., A.C., and A.M.K. V.Z., M.A.J., A.C., and S.S.K. prepared figures for the manuscript.

### Ethics Declarations

V.Z., M.A.J., A.C., A.G., A.M.F., A.M.K., and J.A.N.B. are co-inventors on a filed provisional patent application No. PCT/US24/58838 (USA, 2024) on this work. A.M.K. is a co-founder of the commercial entity Nitro Biosciences. The other authors declare no competing interests.

### Data and material availability

All plasmid materials and bacterial strains will be made available through Addgene or the Bacillus Genetic Stock Center. Sequence files, processed and unprocessed images, and raw data are available as supplementary materials and source data.

