## Supplementary Figures for "Genetic code expansion enables plant-directed control of bacterial activity"

**Supplementary Fig. 1.** Schematic of the co-cultivation luciferase assay using a bacteria-agar matrix to demonstrate bacterial incorporation of plant-produced OMY.

**Supplementary Fig. 2.** Representative scatter plots for flow cytometry gates for data in Extended Data Fig 5. Forward vs side scatter area dot plot with (a) Representative gates drawn on UD1022 after growth in LB and (b) applied to sample of UD1022 reporter extracted from soil after 3 days.

**Supplementary Fig. 3.** Annotations of peak and midpoint locations for luminescence intensity profiles, used to calculate the fold change and max intensity values shown in Fig. 2, Fig. 4, and Extended Data Fig. 2.

**Supplementary Table 1.** List of strains used in this study.

**Supplementary Table 2.** List of constructs used in this study. Full sequences are provided as GenBank files.

**Supplementary Table 3.** List of primers used in construct assembly.

**Supplementary Table 4.** Sample information for proteomics.

**Supplementary Sequences**. GenBank files of constructs used in this study. Genebank files include primers used to construct sequences.

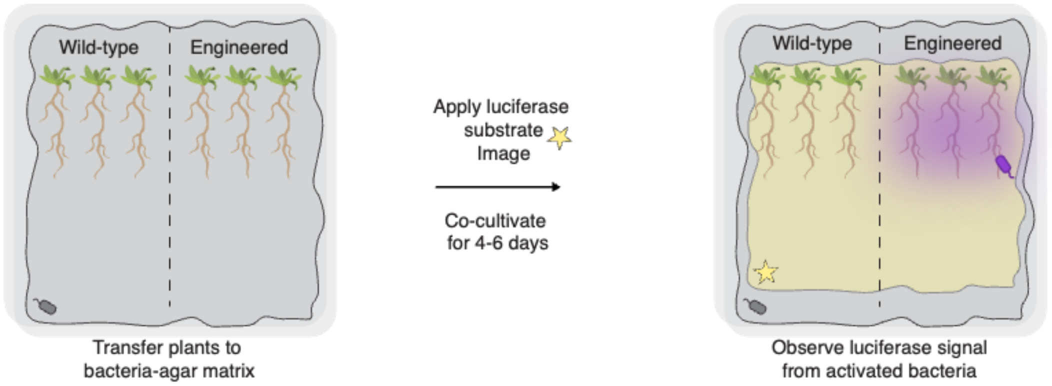


**Supplementary Fig. 1.** Schematic of the co-cultivation luciferase assay using a bacteria-agar matrix to demonstrate bacterial incorporation of plant-produced OMY.

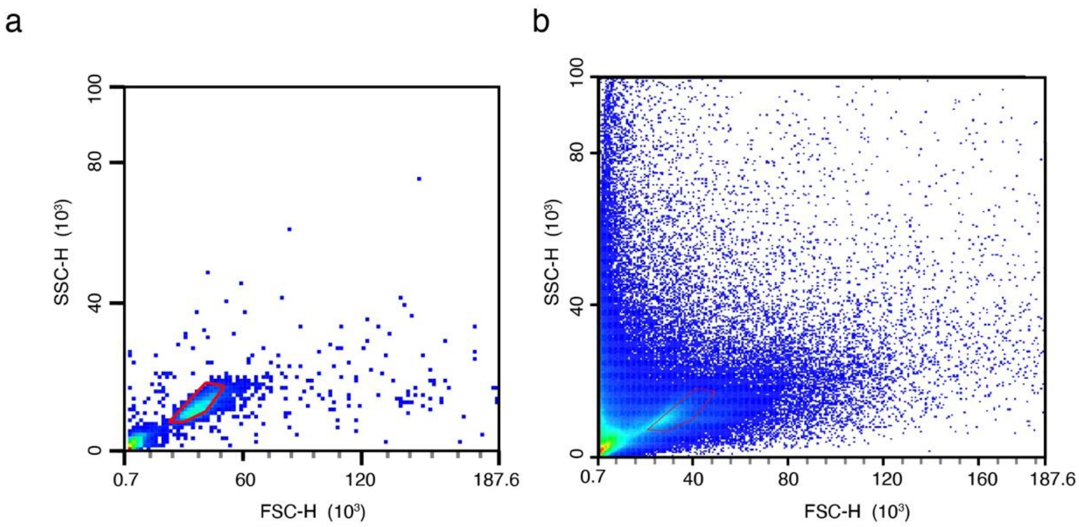


**Supplementary Fig. 2.** Representative scatter plots for flow cytometry gates for data in Extended Data Fig 5. Forward vs side scatter area dot plot with (a) Representative gates drawn on UD1022 after growth in LB and (b) applied to sample of UD1022 reporter extracted from soil after 3 days.

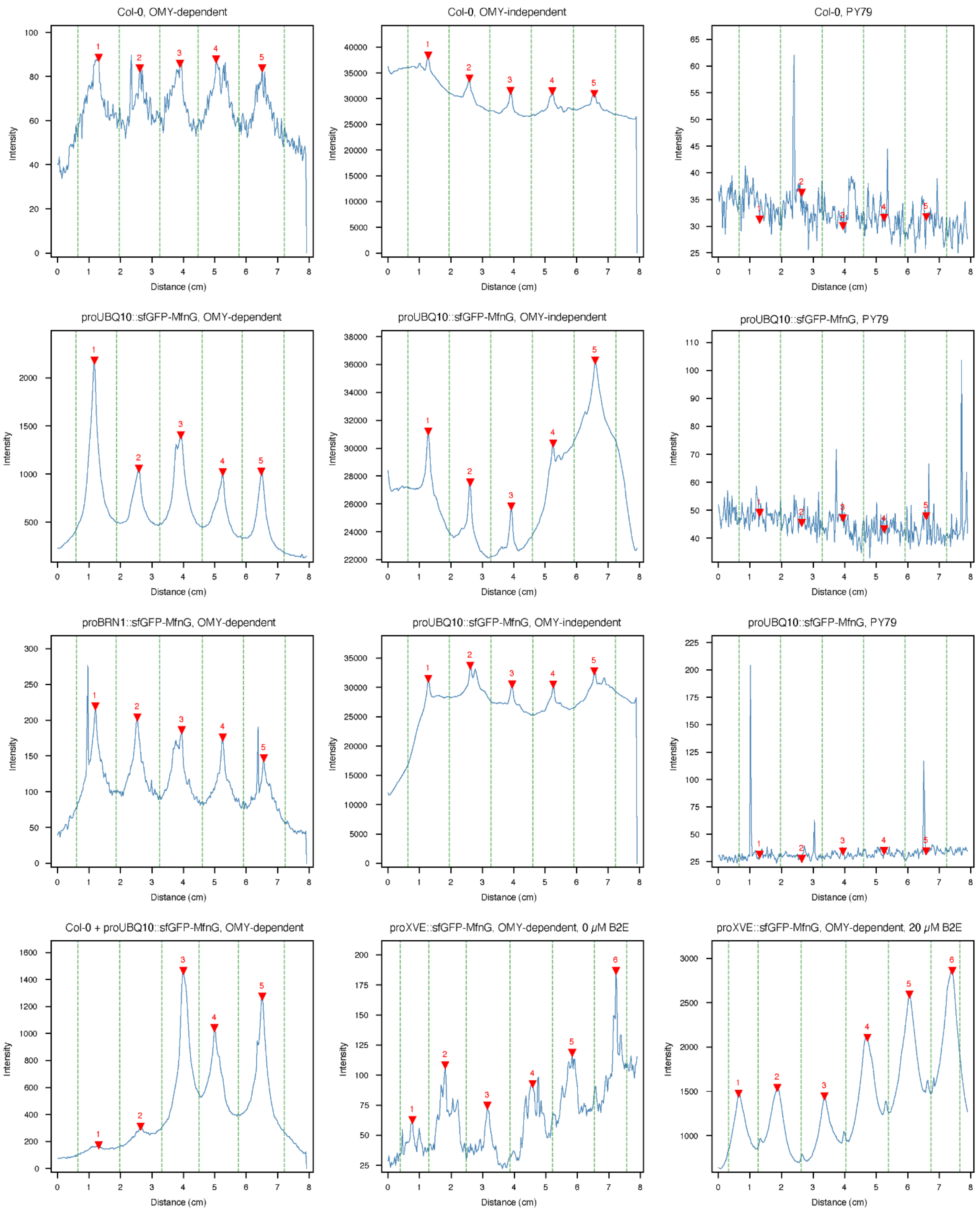
Col-0, OMY-dependent
Col-0, OMY-independent
Col-0, PY79
proUBQ10::sfGFP-MfnG, OMY-dependent
proUBQ10::sfGFP-MfnG, OMY-independent
proUBQ10::sfGFP-MfnG, PY79
proBRN1::sfGFP-MfnG, OMY-dependent
proUBQ10::sfGFP-MfnG, OMY-independent
proUBQ10::sfGFP-MfnG, PY79
Col-0 + proUBQ10::sfGFP-MfnG, OMY-dependent
proXVE::sfGFP-MfnG, OMY-dependent, 0 μM B2E
proXVE::sfGFP-MfnG, OMY-dependent, 20 μM B2E
Intensity
Distance (cm)

**Supplementary Fig. 3.** Annotations of peak (red arrow) and midpoint locations (green dashed line) used to calculate the fold change and max intensity luminescence values shown in Fig. 2, Fig. 4, and Extended Data Fig. 2.
